# *ComPRePS*: Unlocking Scalable AI Analysis for Computational Renal Pathology

**DOI:** 10.1101/2024.03.21.586102

**Authors:** Anindya S. Paul, Luis A. Rodrigues, Suhas Katari Chaluva Kumar, David Manthey, Samuel P. Border, Clara Pardinhas, Carolina Pimenta, Vitor Sousa, Arnaldo Figueiredo, Rui Alves, Briana A. Santo, Will Dunklin, Sumanth Devarasetty, Haitham Abdelazim, Agnes B. Fogo, Avi Z. Rosenberg, Oleksandr Moskalenko, Pinaki Sarder

**Author notes:** Corresponding author: Pinaki Sarder,. These authors contributed equally to this work (Anindya S. Paul, Luis A. Rodrigues).

## Abstract

Digital pathology using whole slide imaging (WSI) and artificial intelligence (AI) has the potential to transform diagnostic workflows, but adoption remains limited by technical complexity and scalability. We developed the Computational Renal Pathology Suite (ComPRePS), a scalable cloud-based platform that automates WSI ingestion, compartmental segmentation, feature extraction, and AI-assisted interpretation through an integrated high-performance architecture.

ComPRePS was evaluated in two use cases. First, using 213 procurement biopsies, we compared conventional assessments with automated AI analyses and a hybrid AI-assisted expert workflow. ComPRePS AI-assisted methods achieved higher precision and significantly improved interobserver agreement for key lesions, including global glomerulosclerosis, interstitial fibrosis and tubular atrophy, and arterial intimal thickening. Second, ComPRePS enabled high-throughput quantitative profiling of glomerular and tubular features across minimal change disease, diabetic nephropathy, and amyloid nephropathy revealing disease-specific phenotypic patterns inaccessible to manual evaluation.

Overall, ComPRePS improves reproducibility, interpretability, and objectivity in renal pathology, bridging computation with clinical practice.

## Introduction

Histopathologic evaluation is essential for diagnosing and managing renal diseases, from chronic kidney conditions to acute transplant rejection, but current workflows remain manual, time-consuming, and prone to variability. Tasks such as lesion counting and fibrosis scoring require expertise yet still yield poor interobserver agreement^1–3^. Manual quantification is also impractical for large-scale studies and requires manual translation of findings to large scale data frames using data collection forms such as spreadsheets^4^. Whole slide imaging (WSI) and artificial intelligence (AI) now offer digitized, scalable approaches that can improve reproducibility and efficiency^5–10^, but clinical adoption is hindered by non-standardized image formats, staining variability, limited interpretability, and steep technical requirements^11^. Open-source platforms such as QuPath^12^ and Automated Slide Analysis Platform (ASAP)^13^ demand advanced knowledge and remain restricted to desktop environments, limiting scalability and collaboration, whereas commercial solutions, though user-friendly, are costly and equally constrained.^14^ These challenges highlight the need for standardized, accessible, and interoperable AI-enabled tools that can bridge research and clinical practice.

In this study, we present ComPRePS, an AI-enabled computational pathology platform that combines a horizontal scalable cloud-HPC infrastructure with a robust, plugin-based architecture built on top of a standardized open-source format. This design enables rapid parallel analysis of hundreds of WSIs through a unified backend architecture that integrates data management, scalable file handling, and distributed task execution. By seamlessly integrating computational and storage resources, the platform allows users to perform large-scale analyses with on-demand access to high-performance infrastructure, effectively enabling complex workflows to be executed at the press of a button, facilitating broad adoption across research and clinical settings. The platform also hosts a diverse “model zoo”, curated collection of pre-trained, standardized models and weights, encompassing internal pipelines for multi-compartment kidney segmentation – including glomeruli, tubules, arteries, arterioles, and arterial intima^15^ – as well as modules for Interstitial Fibrosis and Tubular Atrophy (IFTA) detection^16^, alongside externally developed tools such as peritubular capillary (PTC) segmentation^17^ and DeepCell^18^. Unlike previous systems, these pipelines are actively used for large-scale data generation and analysis across national initiatives and international collaborators^19,20^, underscoring their real-world impact, interoperability, and translational relevance. Beyond segmentation, ComPRePS provides advanced feature extraction and visualization tools that enable automated morphometric analysis across biological, pathomic, and clinical levels, while also offering interactive editing and quality control of the AI results through the user-interface (UI). The platform maintains complete provenance of different user activities on each WSI, enabling transparency and large-scale, multi-institutional collaboration on shared datasets around the world. A major focus of the platform’s design was to enhance user experience and facilitate its integration into daily diagnostic workflows, allowing practical use in routine pathology observations. Users can interactively explore quantitative descriptors, compare features across images, cohorts, or disease groups, and generate dynamic visualizations—such as scatterplots, violin plots, and clustering maps—with direct export of structured results. These capabilities markedly improve accessibility and reproducibility compared with conventional script-based workflows typically required by most open-source tools. The real-world utility of ComPRePS has been demonstrated in large National Institutes of Health (NIH) funded consortia such as the Kidney Precision Medicine Project (KPMP), where leading nephropathologists have used its automated compartmental segmentation, pathomic feature generation, and interactive human-AI quality control workflow to achieve highly accurate, consistent, and reproducible tissue assessments.

To demonstrate the practical impact of ComPRePS in a real-world clinical setting, we designed two illustrative use cases. The first highlights how AI-driven analysis can support pathologists and reduce interobserver variability in evaluating deceased donor procurement biopsies for kidney transplantation. Procurement or harvest biopsies, although frequently used to evaluate marginal donor kidneys worldwide, rely on non-specialist interpretation and leads to inconsistent clinical decisions^21–27^. This variability contributes to high discard rates, further exacerbating the persistent imbalance between kidney transplant supply and demand, where high waitlist mortality coexists with the frequent discard of deceased donor kidneys^28–30^. In the second use case, we applied ComPRePS to quantify morphological differences in renal functional tissue units (FTUs) across three disease groups: Amyloid Nephropathy, Diabetic Nephropathy, and Minimal Change Disease. Manual morphometric analysis of FTUs in large-scale histopathological datasets and correlation with patient clinical data and diagnostic outcome is labor-intensive and needs vast expertise in renal pathology, whereas computational tools enable consistent and scalable analyses and exhibit the potential to standardize the mapping between biopsy findings with diagnostic results. Accessible and explainable approaches for integrating pathomic features into clinical decision-making - from clinically interpretable metrics such as glomerular size, area, and mesangial fraction to complex image morphometrics including nuclear contrast, texture, shape, and size - could transform digital pathology from a descriptive to a dependable predictive discipline, enabling disease mechanism discovery and outcome prediction.

ComPRePS has been operationally adopted within the KPMP for data generation in the NIH-funded Kidney Tissue Atlas initiative, a multi-year effort to advance mechanistic understanding of kidney disease. Within KPMP program, ComPRePS has been used to generate high-quality, standardized pathomic derivatives from patient biopsies, now integrated into the publicly accessible KPMP Tissue Atlas^20^. This integration underscores the platform’s ease-of-use, translational maturity and its contribution to open, data-driven kidney research.

## Results

### Collaborative Workflows in ComPRePS

ComPRePS was designed as a collaborative platform where diverse user groups from across the world can bring their expertise to advance AI research in the medical field. Figure 1 illustrates the schematic workflow showing the three main stakeholders within the ecosystem - developers, researchers, and clinicians. Developers contribute by developing algorithms and integrating their AI workflows into ComPRePS that can be executed reproducibly on a large-scale. Researchers provide domain expertise, curate datasets, harmonize various diseased cohorts, generate annotations, and define clinically relevant problems that drive model development. Clinicians use the AI models resulting from developer-researchers’ collaboration through the ComPRePS interface to obtain insights that directly inform decision-making in diagnostic or research settings.

**Figure 1.**
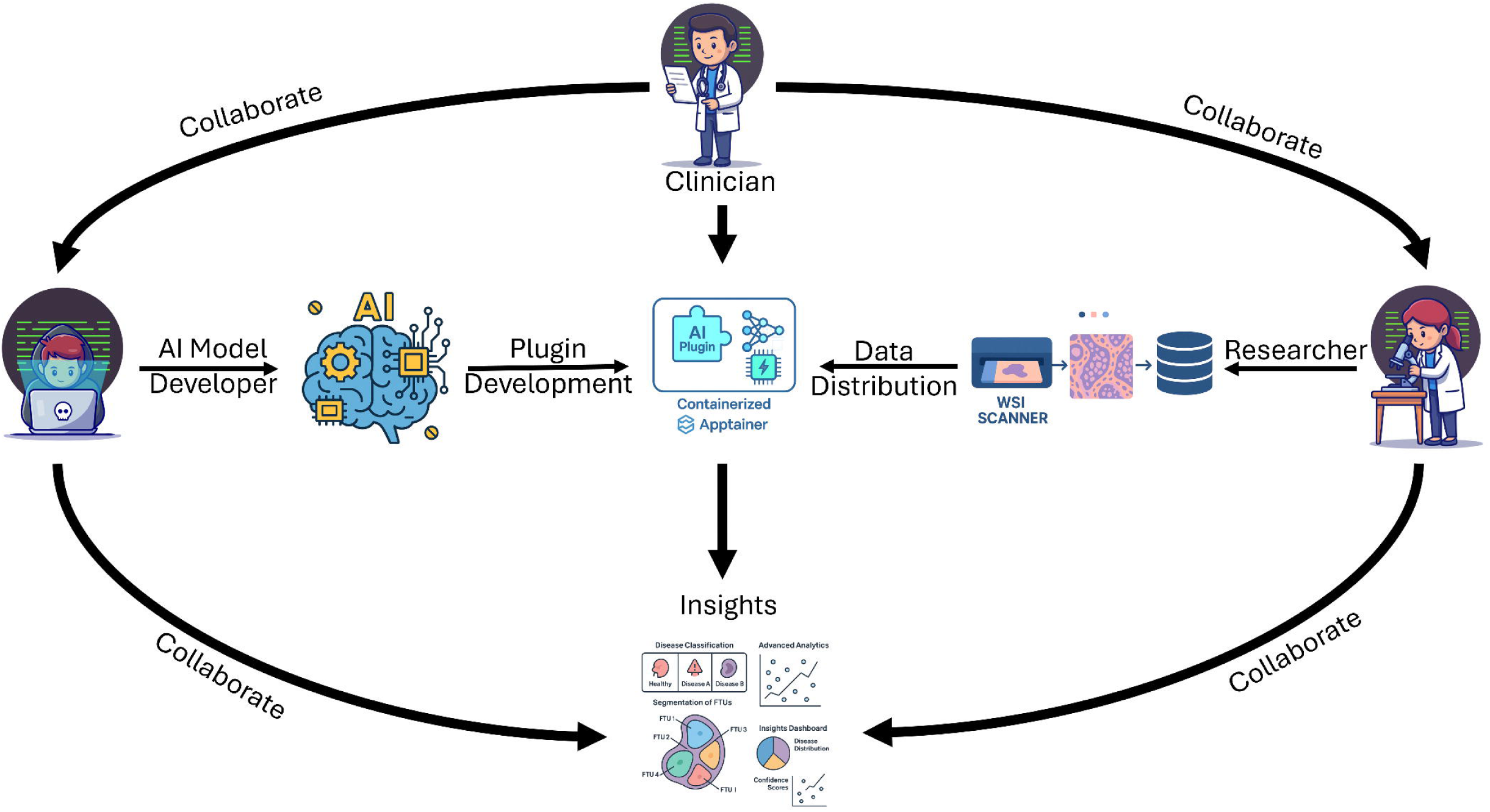
Schematic Workflow diagram describing how developers, researchers and clinicians work in collaboration to perform image analysis using ComPRePS.

This workflow creates a feedback loop in which clinical priorities guide algorithm development, research validation strengthens model credibility, and end users benefit from streamlined access to advanced tools. The modular architecture and ComPRePS’s tools allow these groups to collaborate seamlessly without requiring in-depth knowledge of how to deploy AI in a massive scalable architecture-processing efficiently on giga-pixel sized WSIs.

#### Global Hub for Collaboration

Figure 2 demonstrates one of the most powerful and novel capabilities of ComPRePS: the ability to act as a global hub for model dissemination. Consider a scenario where an AI developer, working anywhere in the world, identifies a pressing problem in the medical domain. Using publicly available datasets such as those provided by KPMP or the HuBMAP consortium or their own private datasets, they can train and refine their model in close collaboration with pathologists, researchers and clinicians from any institution across the world. Once the developers are satisfied with the performance of the model, the next step is for them to package it as a plugin in the Slicer-CLI format and containerized as a Docker Image, ready for deployment. This example demonstrates the potential of a novel partnership between multidisciplinary experts to harmonize cutting-edge AI tools with real-world clinical data, setting a new standard for precision medicine in kidney research and diagnostic applications.

**Figure 2.**
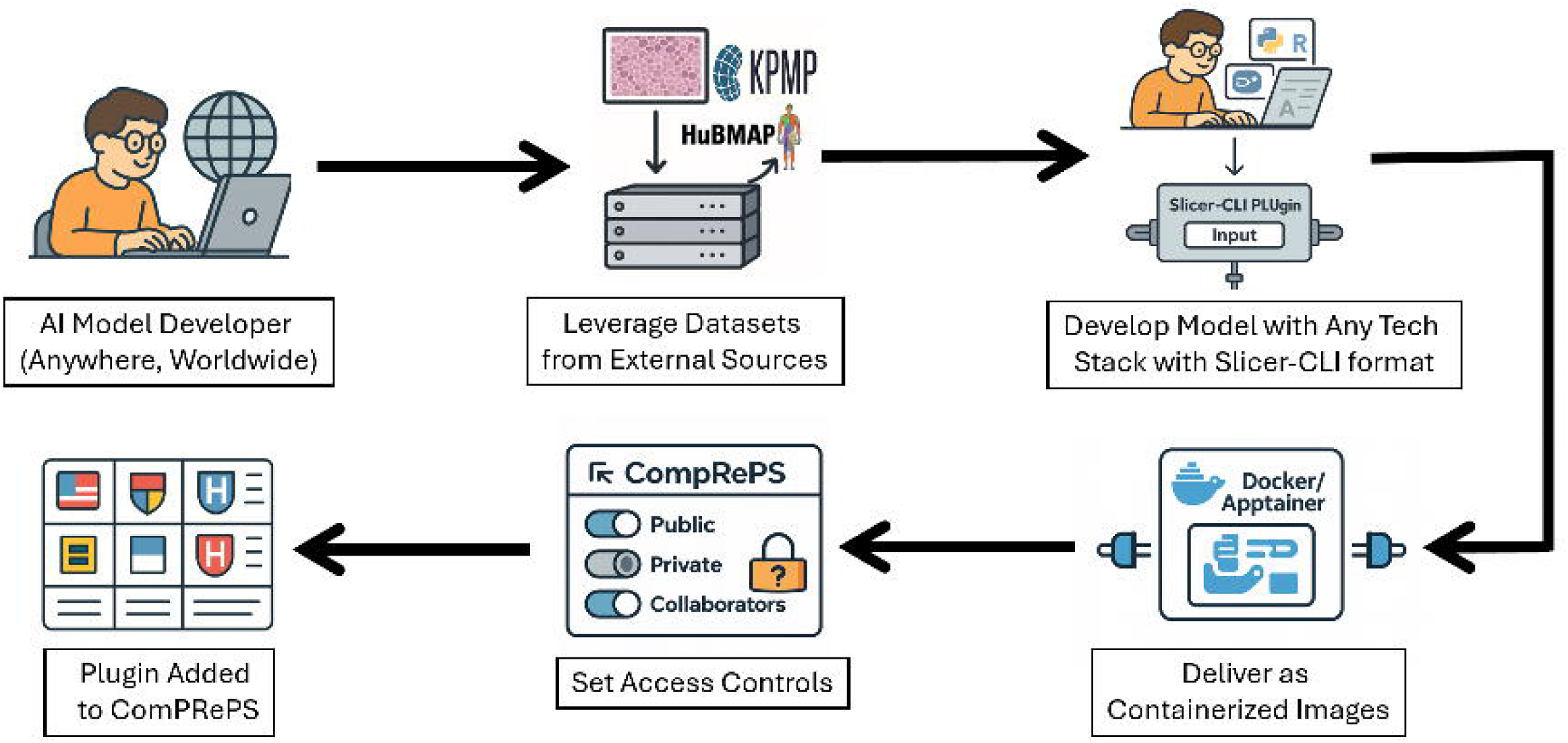
Schematic Workflow diagram describing how AI Model Developers from across the world can tackle challenging medical problems with ComPRePS. Abbreviations: AI, artificial intelligence; ComPRePS, Computational Renal Pathology Suite; KPMP, Kidney Precision Medicine Project; HuBMAP, Human Biomolecular Atlas Program; Slicer-CLI, 3D Slicer Command Line Interface.

At this point, ComPRePS becomes the bridge between innovation and real-world impact. The developer can seamlessly deploy their model within ComPRePS, where it can be tested, validated, and shared with the broader medical and scientific research community. Flexible access controls allow them to collaborate privately with specific groups, or, alternatively, they can disseminate the tool widely to accelerate adoption and gather invaluable feedback. In this way, ComPRePS functions not only as a scalable AI compute platform, but also as a global hub for scientific collaboration, enabling developers to move from model conception to community driven application with unprecedented efficiency.

Importantly, the open-source nature of the platform ensures that tools are not limited to select individuals and institutions, but can be deployed broadly, accelerating reproducibility and enabling large-scale validation across diverse populations.

In addition, ComPRePS Global Hub can act as a unified model repository in digital pathology for accelerating translational research and democratizing access to advanced AI tools in the medical community. By consolidating diverse, well-validated models developed worldwide - ranging from tissue segmentation and cellular quantification to feature extraction and outcome prediction - within a unified, interoperable framework, ComPRePS model zoo enables researchers and clinicians to rapidly validate and benchmark algorithms across varied datasets and disease contexts. As pathological evaluation of tissue images in clinical pipeline have demonstrated significant intra and inter observer variability, a standardization and shared ecosystem of development and deployment enhance reproducibility, reduces technical point-of-entry of adopting AI and finally foster inter-institution collaborative learning.

#### Scalability and Adoption

Figure 3 highlights ComPRePS’s user-friendly interface and how users can easily visualize the results of AI segmentations as an overlay on the original WSI. The interface enables clinicians to toggle between annotation layers, view segmentations at multiple magnifications, and explore extracted features through interactive plots via the plotting tool. For researchers, the ability to aggregate and compare features across datasets facilitates hypothesis generation and cohort-level analyses. For developers, the clear visualization of output supports rapid evaluation of model performance. These features establish ComPRePS as not only a computational platform but also a translational interface where outputs can be directly interpreted and applied in biomedical contexts. An ongoing usability study is further evaluating how the visualization interface supports workflow efficiency, interpretability, and adoption across multidisciplinary users.

**Figure 3.**
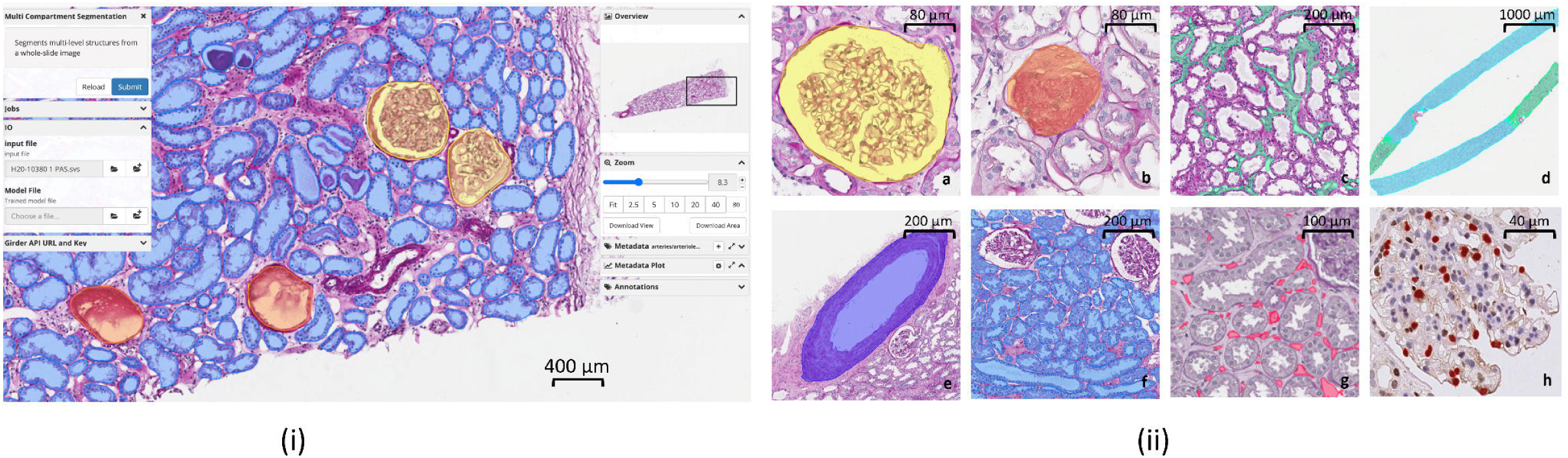
Overview of ComPRePS UI and examples of AI-generated segmentation of renal FTUs. (i) ComPRePS interface showing segmented FTUs: Non-Globally Sclerotic Glomeruli (yellow), Globally Sclerotic Glomeruli (red), and Tubules (blue) (scale bar: 400 µm). (ii) High-magnification AI-generated FTU segmentations with corresponding scale bars: a—Non-Globally Sclerotic Glomerulus (scale bar: 80 µm); b—Globally Sclerotic Glomerulus (scale bar: 80 µm); c—Cortical interstitial space (scale bar: 200 µm); d—Cortical (green) and medullary (blue) regions (scale bar: 1000 µm); e—Artery (scale bar: 200 µm); f—Cortical tubules (scale bar: 200 µm); g—Peritubular capillaries (scale bar: 100 µm); h—Podocyte nuclei (scale bar: 40 µm).

The adoption of ComPRePS by consortia and academic partners demonstrates its scalability and practical value. Pipelines such as Multi-Compartment Segmentation and IFTA Segmentation are routinely used by KPMP, HuBMAP, Indiana University, and the University of Coimbra, with AI-outputs contributing to the generation of large open-source data Atlas^19,20^. We have contributed AI-segmented kidney compartments along with detailed morphometric feature data from over 400 tissue slides to the KPMP 2.0 Atlas, enhancing ComPRePS’s reputation as a large-scale data generation platform for the open-science community. Collaborative efforts have already led to the development of new pipelines, such as the arterial intima Segmentation model created in collaboration with the University of Coimbra.

#### Interoperability and External Integration

Beyond its native plugins, ComPRePS has been adopted by external platforms as a computational service. For example, the recently published multi-modal integration tool between spatial OMICS and histological morphometrics: FUSION^31^ application uses ComPRePS’s APIs for high-throughput AI processing as well as availability of datasets. This interoperability demonstrates the flexibility of the platform, potential of rapid tool development in serving beyond histopathology, and its ability to support heterogeneous workflows without requiring end users to manage infrastructure.

Such integrations with histopathological morphometrics and molecular profiles are critical for building connected ecosystems where AI pipelines, visualization platforms, and clinical decision-support tools can interact seamlessly across domains. By exposing standardized interfaces, ComPRePS lowers the barriers for adoption and promotes the reuse of validated pipelines in diverse contexts.

### Clinical Use - Case 1: Donor biopsy assessment for kidney transplant

#### AI Augmented Biopsy Evaluation

The ComPRePS user-interface enabled reviewing pathologists to assess each WSI with the support of automated AI-augmented segmentation and lesion scores. Figure 4 displays this interface. For glomerulosclerosis, all glomeruli (sclerotic and non-sclerotic) were visualized, along with the computed percentage of sclerotic glomeruli and the corresponding lesion score (Figure 4a). For IFTA, the plugin segmented fibrotic and atrophic tubular regions quantified as a percentage of the cortical area (Figures 4b and 4c). For arterial intimal fibrosis, Figure 5 illustrates ComPRePS-generated fibro-intima predictions and stenosis ratios across multiple arterial orientations and varying degrees of thickening. This allowed the pathologist to select the arterial orientation that best corresponded to the tissue section, which was particularly useful for arteries in longitudinal sections.

**Figure 4.**
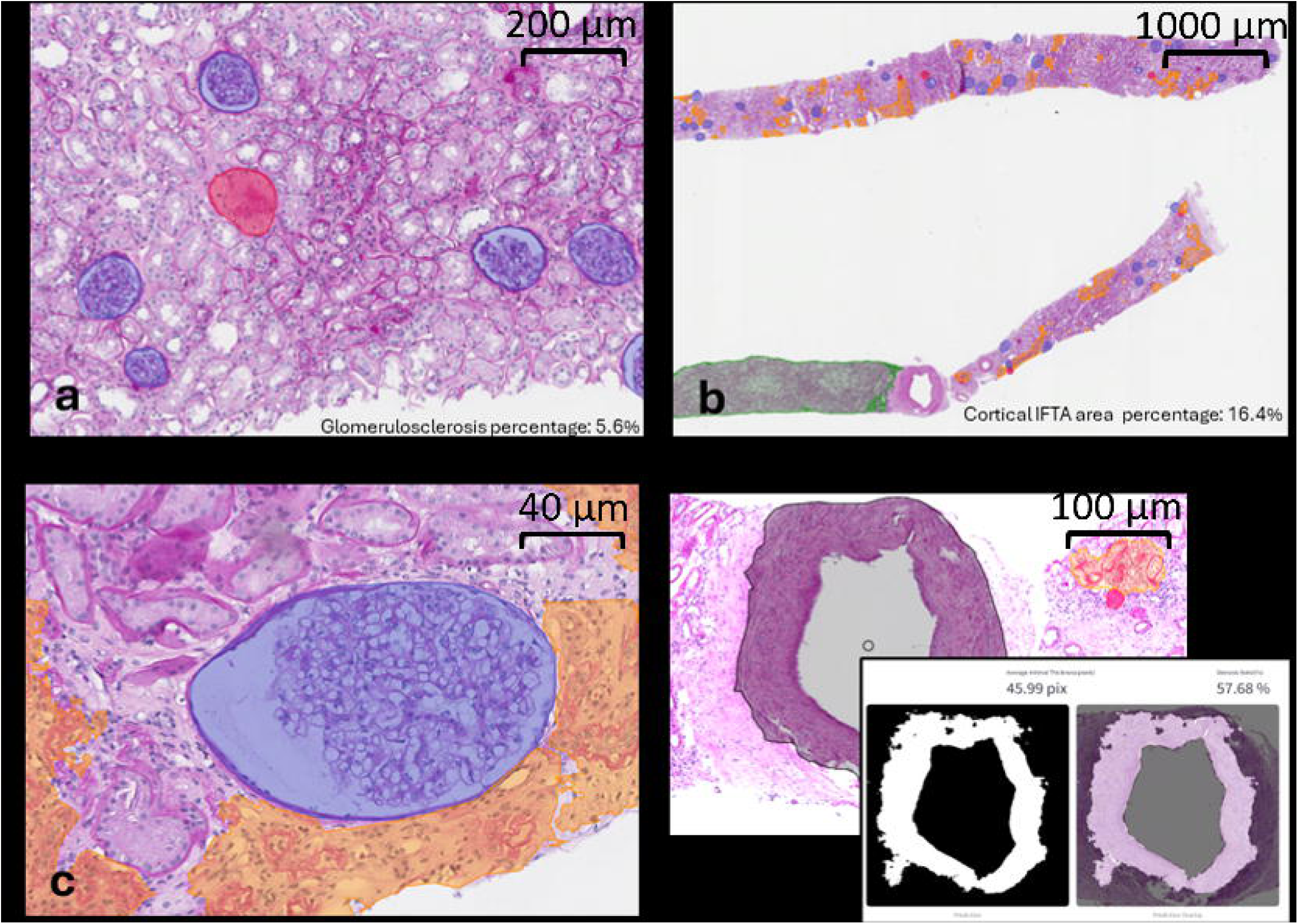
ComPRePS human–AI assisted evaluation of the Remuzzi Score in procurement kidney transplant biopsies. Example case with total Remuzzi score = 6: glomerulosclerosis < 20% (score = 1), IFTA < 20% (score = 2), and arterial intimal thickening > 50% (score = 3). (a) Glomerular segmentation showing globally sclerotic (red) and non-sclerotic (blue) glomeruli with interactive validation and automated percentage output (scale bar: 200 µm). (b) AI-generated IFTA overlays enabling rapid screening, QC correction, and cortical IFTA quantification (scale bar: 1000 µm). (c) High-power peri-glomerular IFTA visualization (scale bar: 40 µm). (d) Arterial intimal thickening module: user-selected arteries with automated inner/outer boundary detection (black), IFTA region (orange), and binary mask/overlay outputs (scale bar: 100 µm). Quantitative metrics such as mean intimal thickness and luminal stenosis are displayed.

**Figure 5.**
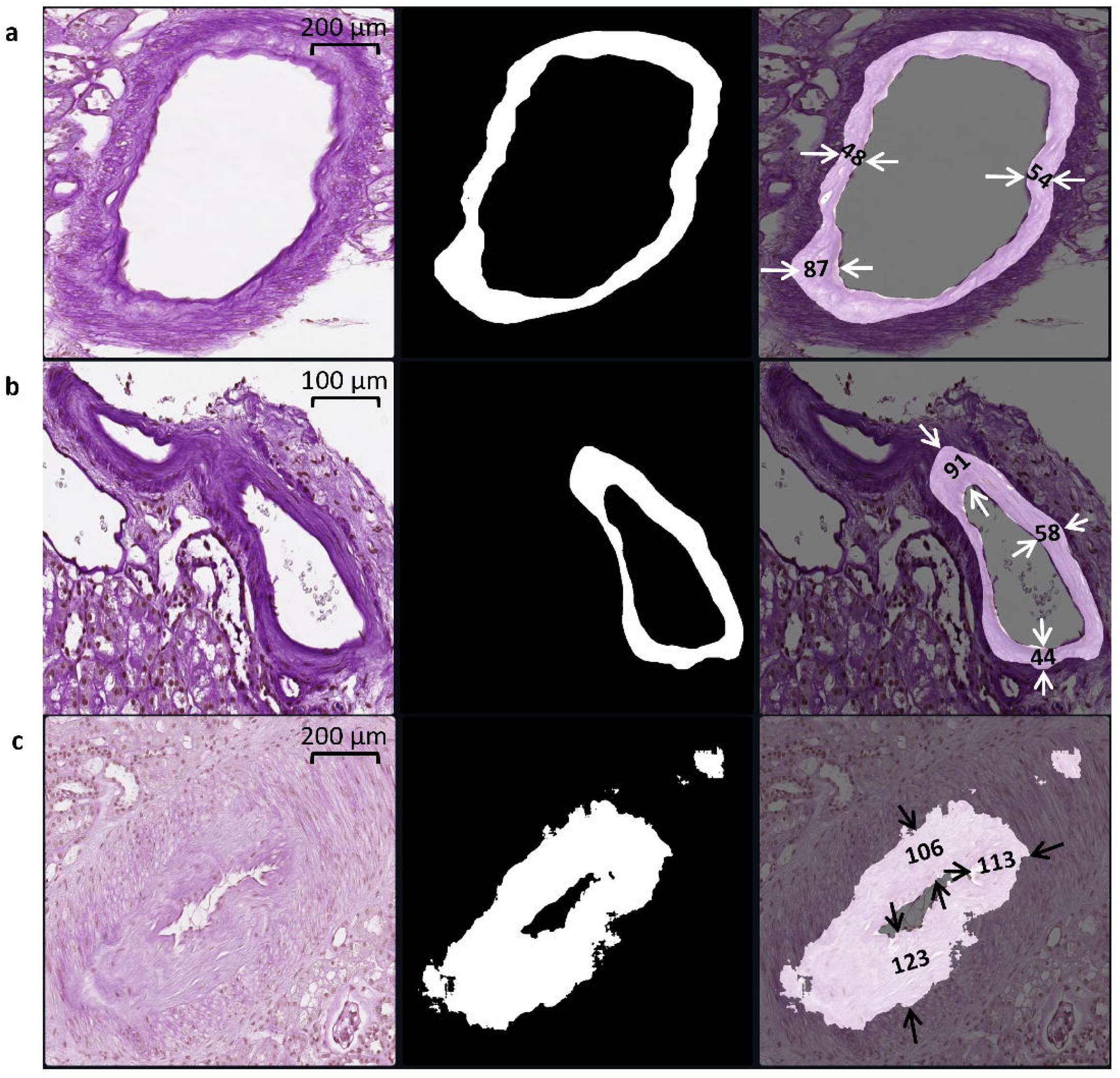
Representative examples of arterial intimal segmentation and fibro intimal lesion quantification, illustrating the model’s performance across different anatomical orientations and degrees of intimal thickening. Each row shows (left to right): the original cropped artery image, the prediction mask, and the overlay with the predicted intimal thickness measurements (arrows, in pixels). Predicted luminal stenosis ratios: (a) 23.3%, grade 1 (scale bar: 200 µm) (b) 35.5%, grade 2 (scale bar: 100 µm) (c) 94%, grade 3 (scale bar: 200 µm). This figure demonstrates how measurement site selection can influence grading outcomes, particularly in longitudinally sectioned arteries.

#### Interobserver Agreement in Remuzzi Scoring

For benchmarking between different biopsy evaluation workflows, we used the reviewer–ComPRePS workflow as the GT due to its reduced interobserver variability and integration of AI-driven precision with expert interpretation.

A comprehensive analysis of interobserver agreement for each Remuzzi score component and the overall score is provided in the *Supplementary Information* (Supplementary Tables 1–7). This includes confusion matrices comparing ComPRePS predictions with GT and GT with on-call pathologist assessments, as well as prediction metrics (precision, recall, F1 score, and accuracy) for ComPRePS relative to on-call pathologists, using GT as the reference standard.

A summary of the overall results for each Remuzzi component and the final score is presented below. Table 1 illustrates the histopathological classes of chronic lesions scored as per Remuzzi classification. Supplementary Table 8 illustrates the final clinical decision making by the on-call pathologists with the corresponding kidney utilization decisions for the donor organ. Table 2 presents Cohen’s Kappa values for pairwise comparisons between pathologists and ComPRePS for glomerulosclerosis, IFTA, and arterial intimal thickening, while Table 3 shows the results for the final Remuzzi score in organ transplant decision-making. Supplementary Table 9 lists the reference intervals for interpreting Cohen’s Kappa values.

**Table 1:** Remuzzi protocol: Semiquantitative method of scoring histopathologic lesions.

|  | 0 | 1 | 2 | 3 |
| --- | --- | --- | --- | --- |
| Global glomerulosclerosis | None | < 20% | 20 - 50% | > 50% |
| Tubular atrophy | Absent | < 20% | 20 - 50% | > 50% |
| Interstitial fibrosis | Absent | < 20% | 20 - 50% | > 50% |
| Arterial intimal fibrosis | Absent | < 25% | 25 - 50% | > 50% |

**Table 2:** From (top to bottom) Cortical IFTA, Glomerulosclerosis, and Arterial intimal fibrosis Cohen’s Kappa values for pairwise comparisons between 4 workflows by 2 expert pathologists: GT: human-AI (gold standard), RP: expert renal pathologist, OP: On-call pathologist, ComPRePS: AI alone.

|  | GT1 | RP1 | GT2 | RP2 | OP | ComPrePS |
| --- | --- | --- | --- | --- | --- | --- |
| GT1 | 1 | 0.61 | 0.57 | 0.42 | 0.23 | 0.56 |
| RP1 |  | 1 | 0.37 | 0.29 | 0.48 | 0.28 |
| GT2 |  |  | 1 | 0.57 | 0.14 | 0.62 |
| RP2 |  |  |  | 1 | 0.13 | 0.31 |
| OP |  |  |  |  | 1 | 0.07 |
| GT1 | 1 | 0.85 | 0.85 | 0.80 | 0.69 | 0.90 |
| RP1 |  | 1 | 0.80 | 0.76 | 0.70 | 0.80 |
| GT2 |  |  | 1 | 0.87 | 0.65 | 0.88 |
| RP2 |  |  |  | 1 | 0.63 | 0.77 |
| OP |  |  |  |  | 1 | 0.63 |
| GT1 | 1 | 0.79 | 0.67 | 0.42 | 0.44 | 0.89 |
| RP1 |  | 1 | 0.56 | 0.47 | 0.55 | 0.72 |
| GT2 |  |  | 1 | 0.49 | 0.36 | 0.86 |
| RP2 |  |  |  | 1 | 0.33 | 0.45 |
| OP |  |  |  |  | 1 | 0.44 |

**Table 3:** Final Remuzzi Cohen’s Kappa values for pairwise comparisons between 4 workflows by 2 expert pathologists: GT: human-AI (gold standard), RP: expert renal pathologist, OP: On-call pathologist, ComPRePS: AI alone.

|  | GT1 | RP1 | GT2 | RP2 | OP | ComPrePS |
| --- | --- | --- | --- | --- | --- | --- |
| GT1 | 1 | 0.73 | 0.69 | 0.47 | 0.42 | 0.69 |
| RP1 |  | 1 | 0.56 | 0.46 | 0.52 | 0.55 |
| GT2 |  |  | 1 | 0.59 | 0.35 | 0.75 |
| RP2 |  |  |  | 1 | 0.31 | 0.48 |
| OP |  |  |  |  | 1 | 0.32 |

#### Glomerulosclerosis

We observed that the comparison between GT and ComPRePS reveals stronger and more consistent performance, particularly in precision, recall, and F1 scores, with notable improvements in challenging classes such as Class 2. GT vs. on-call (OP) shows significantly lower precision and recall, with the most pronounced gap observed in Class 2 recall performance (Supplementary Table 2: 0.90 vs. 0.53). Class 3 results were not considered reliable due to the rarity of donor samples with >50% sclerosed glomeruli (*n* = 2). Intra-class correlation (Table 2) shows that agreement between reviewers – 1 and 2 using AI-generated masks (GT1 vs. GT2) is approximately 0.1 higher than without AI (RP1 vs. RP2), indicating reduced interobserver variability. Moreover, GT1 and GT2 exhibit Kappa values 0.3–0.5 higher with ComPRePS than with each other, and ComPRePS shows Kappa values 1.0–2.0 higher with expert reviewers (RP1 and RP2) compared to on-call pathologists (OP). Cohen’s Kappa values for ComPRePS vs. expert reviewer comparisons mostly fall within the good–excellent range, whereas on-call pathologist agreement remains moderate. These findings highlight the potential of AI integration and pathologist-AI collaboration to enhance lesion classification accuracy, improve expert consensus, and outperform non-specialist assessments in renal pathology.

#### Cortical IFTA

ComPRePS showed better agreement with GT1 and GT2 and stronger classification performance than on-call assessments, especially in Classes 0 and 2. While on-call pathologists slightly outperformed ComPRePS in Class 4, the result was misleading due to class imbalance. As shown in Table 2, the interobserver agreement for cortical IFTA between two expert renal pathologists (RP1 vs. RP2) was low (κ = 0.294) but improved to a moderate level (κ = 0.573) when they used AI-generated segmentation masks (GT1 vs. GT2). To understand this improvement more deeply, we observed each pathologist’s workflow when they used AI-generated IFTA masks compared to when they did not. Pathologists frequently spent considerable time navigating between low to high power magnifications to identify IFTA regions, leading to missed or inconsistent annotations and increased interobserver variability. The use of adjustable AI-generated masks improved both efficiency and accuracy, resulting in markedly reduced variability among observers. On-call pathologists (OP) did not have the benefit of using AI masks and showed consistently low agreement with all other assessors.

##### Arterial Intimal Fibrosis

Overall, classification accuracy for both ComPRePS and pathologists was higher in arteries exhibiting greater intimal thickening and moderate-to-severe chronic injury (Supplementary Tables 5a and 6a). While on-call pathologists and expert reviewers agreed perfectly on normal vessels, accuracy of on-call pathologists declined sharply for arteries graded 1–3, with only about one-third of class 2 and 3 lesions correctly classified. Agreement between two expert renal pathologists improved from low (κ = 0.46) to moderate (κ = 0.67) when using ComPRePS-generated segmentation masks, and ComPRePS incorporating reviewer-defined measurement points achieved high agreement with GT1 (κ = 0.90) and GT2 (κ = 0.86). These findings highlight how a well-designed, interactive human–AI platform can reduce interobserver variability and make vascular lesion scoring in procurement biopsies more quantitative.

#### Final Remuzzi Score

ComPRePS outperformed on-call assessments in predicting the final grades used for organ allocation decisions as shown in Table 3. In critical affirmative categories (e.g., Class 1 for allocation or Classes 6–8 for rejection), ComPRePS achieved significantly higher precision, recall, and F1-scores (Supplementary Table 7) and maintained consistent accuracy across all categories, unlike on-call assessments, which were more variable and less reliable. As shown in Table 3, independent assessments by both expert reviewers and on-call pathologists demonstrated agreement levels below 50%. In contrast, when reviewers partnered with ComPRePS, inter-reviewer agreement between Reviewer 1 and 2 was substantially improved (from κ score 0.46 to 0.69). The results underscore the potential of AI-assisted biopsy evaluation to enhance the precision and consistency of organ allocation decisions, thereby improving organ utilization efficiency and significantly reducing discard rates.

### Clinical Use-case 2: Quantitative Morphological Analysis for Kidney Diseases: A study in AN, DN and MCD cohort

With ComPRePS, we quantified morphological differences in segmented FTUs, specifically glomeruli and tubules, across three disease groups. By extracting and analyzing quantitative features in scale across individual slides and diseases, ComPRePS revealed group-specific patterns in tissue architecture, capturing both subtle and overt structural alterations. The results of these analyses are described below.

### Glomerular Analysis

#### Morphological analysis of glomerulosclerosis present in a WSI

After running our multi-compartment segmentation tool for segmenting glomeruli, ComPRePS offers a range of feature extraction methods to automate kidney histomorphometry measurements at multiple levels of granularity. To analyze glomerulosclerosis within a single WSI, we focused on the quantification of the eosinophilic area in glomeruli. Figure 6 i-d shows the distribution of the eosinophilic area, normalized to total glomerular area, which is skewed toward higher values. The median eosinophilic content was 75%, with a slightly lower mean of 73%, indicating mild skewness. A chi-square goodness-of-fit test (*p* = 0.03) confirmed a significant deviation from normality supporting the presence of skewness.

**Figure 6.**
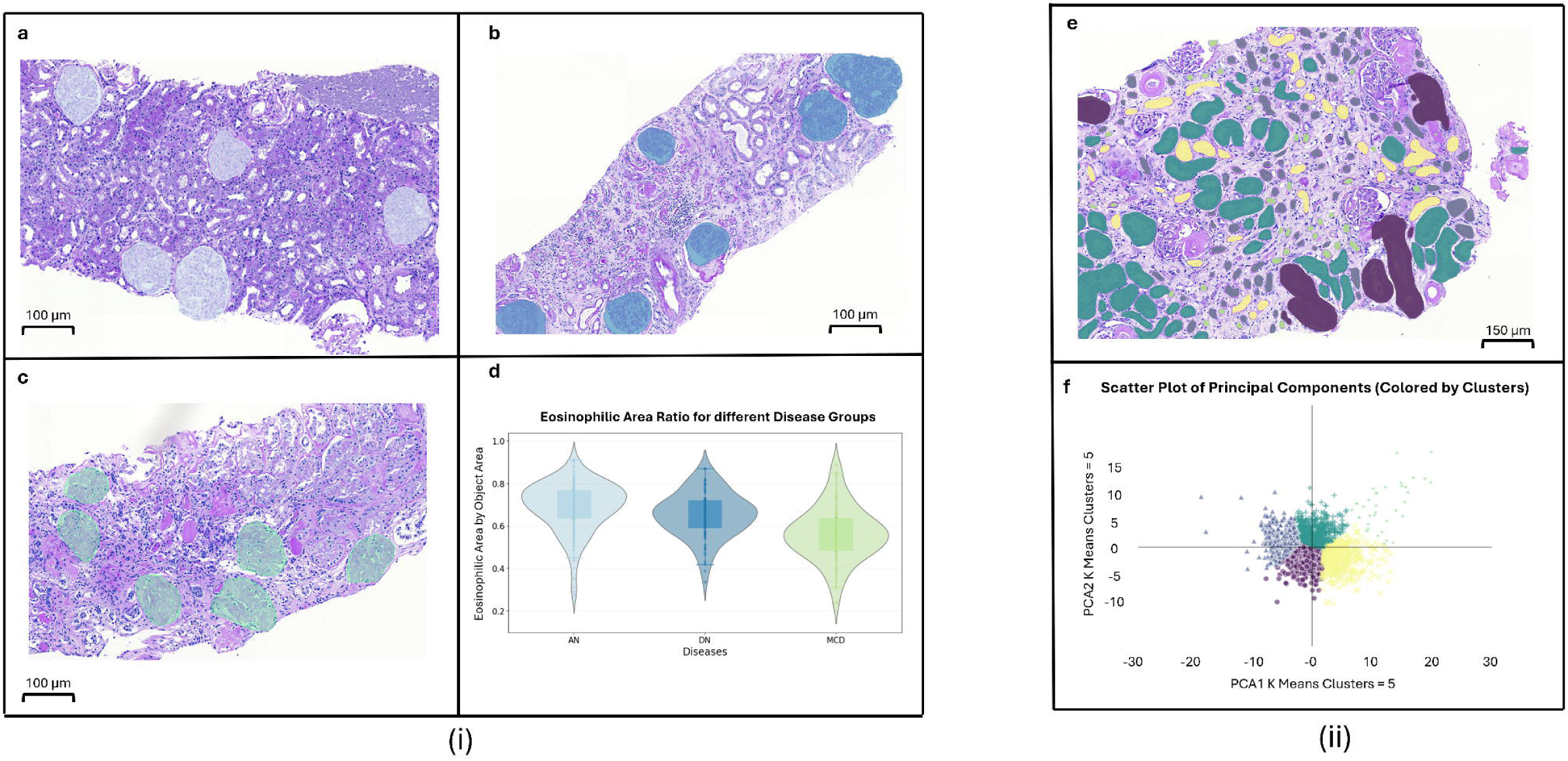
Visualization of quantitative analysis of glomeruli and tubules across different renal disease groups. (i) Segmented glomeruli across representative disease slides: (a) Amyloidosis Nephropathy (AN) (scale bar: 100 µm), (b) Diabetic Nephropathy (DN) (scale bar: 100 µm), and (c) Minimal Change Disease (MCD) (scale bar: 100 µm). (d) Violin plot of eosinophilic area normalized by glomerular area across the three disease groups. (ii) Scatter plot of PCA analysis with 5 clusters run on segmented tubules for an AN case (f), where clusters are color-coded and shown on histology for spatial correlation and visualization (e) (scale bar: 150 µm).

ComPRePS also enables interactive selection of specific ranges within the metadata plot. For instance, filtering above the third quartile isolates the most sclerotic glomeruli—borderline globally sclerotic—which are noticeably smaller than their non-sclerotic counterparts. This finding is consistent with established evidence that glomerular size decreases following marked mesangial expansion and sclerosis^32,33^.

A scatter plot (shown in Supplementary Figure 5a) of eosinophilic area as a fraction of object area on the y-axis with the total object area on the x-axis, where each point corresponds to a glomerulus, illustrated a strong negative correlation between these two morphological features across all glomeruli within this slide. A linear regression analysis yielded an *R*^2^ value of 0.94, confirming that the eosinophilic area has a well-defined inverse relationship with the total object area. This finding corroborates the established pathological observation that increasing glomerulosclerosis - characterized by reduced cellularity and expansion of eosinophilic regions - is associated with a corresponding decrease in glomerular size. Using ComPRePS’s advanced morphometric visualization capabilities, we also uncovered a strong inverse relationship between the mesangial area and capillary area (Supplementary Figure 5b). ComPRePS morphometrics and insights are summarized from its visualization tool after examining every glomerulus and supports the established convention that mesangial expansion, resulting from increased extracellular matrix deposition and mesangial cell proliferation, compresses the glomerular capillary lumen, thereby reducing the effective filtration surface and contributing to the development of glomerular hyperfiltration in proteinuric diseases^34^. This is a simple experiment to demonstrate that by enabling large-scale, quantitative analysis across thousands of biopsies, ComPRePS possesses an unprecedented capability to affirm, refine, or even challenge established pathological paradigms – delivering scale, reproducibility and objectivity far beyond the limits of standard human visual assessment.

#### Comparing glomerular morphology across WSIs and diseases

ComPRePS analyses revealed distinct, disease-specific glomerular patterns that are consistent with the underlying pathological impact. DN showed mesangial expansion with Kimmelstiel-Wilson nodules, AN demonstrated protein aggregation, and MCD appeared largely unremarkable. Figure 6 i-d, violin plot of eosinophilic area relative to object area, demonstrated clear separation among groups, with AN and DN glomeruli containing ∼20% more eosinophilic tissue than MCD.

Quantitative texture analysis further distinguished groups: eosinophilic regions in AN and DN were smoother and more homogeneous than those in MCD, with values lowest in AN, intermediate in DN, and highest in MCD (AN < DN < MCD) (Figure 6 i-d).

In addition to exhibiting a greater eosinophilic area, AN and DN glomeruli displayed distinct textural features compared to MCD. Texture was quantified using the gray-level co-occurrence matrix (GLCM), with energy selected as the representative metric. Lower energy values indicate smoother and more homogeneous regions, whereas higher values reflect greater intensity variation. Consistent with this, energy values followed the pattern AN < DN < MCD. A one-way ANOVA confirmed significant differences across groups (*p* = 1.82 × 10⁻¹⁵). Post-hoc Tukey HSD analysis demonstrated significantly lower energy in AN versus MCD (*p* < 0.0001), DN versus MCD (*p* < 0.0001), and AN versus DN (*p* = 0.0172). These results indicate that AN and DN glomeruli contain smoother, more homogeneous eosinophilic regions compared to MCD. By performing fine-grained textural and spatial tissue analysis, ComPRePS revealed subtle patterns invisible to human vision in differentiating various glomerular diseases, expanding the scope of human-AI partnership of pathological diagnostic and prognostic interpretation.

### Tubular Analysis

#### Clustering Tubules and Identifying Key Differential Characteristics

Figure 6 ii-f presents a principal component analysis (PCA) of tubular features from the AN cohort (*n* = 5 clusters). The primary differences across clusters were driven by total tubular area and major axis length. Together, these parameters reflect both sectioning angle (“cut”) and tubule type. Smaller tubules were more likely medullary (e.g., descending or ascending thin limbs), whereas larger tubules were typically cortical (proximal or distal). In the PCA plot, clusters varied from small to large (left to right) and long to short (top to bottom). This demonstrated that cluster analysis in high dimensional structural morphometrics beyond human capability using advanced ComPRePS visualization tool reveals spatial heterogeneity and quantifies disease specific tubular alterations as disease progresses, which can broaden our scientific understanding of its impact in long-term kidney function decline.

## Discussion

In this article, we presented ComPRePS, a cloud-based platform designed for kidney data (WSI and clinical metadata) visualization, segmentation, quality control, AI-driven analytics, seamless collaboration, and data management. A core objective of ComPRePS is to democratize AI by adhering to the FAIR (Findable, Accessible, Interoperable, and Reusable) principles outlined by the National Institutes of Health (NIH). By bridging the gap between research and clinical application, ComPRePS aims to address several longstanding challenges in AI adoption, including the elimination of technical barriers to AI tool usage, real-time collaboration among pathologists, and cloud-first accessibility. The platform enables the seamless replication of computational pipeline results with minimal user intervention, fostering reproducibility, and efficiency. The AI and feature extraction tools integrated within ComPRePS are actively utilized in real-world applications, demonstrating their effectiveness in large-scale biomedical research. Specifically, ComPRePS plugins serve as the foundation for analyzing samples provided by multiple institutions across the United States as part of major National Institutes of Health (NIH) consortia, including the Kidney Precision Medicine Project (KPMP)^20^ and the Human Biomolecular Atlas Program (HuBMAP)^19^. In KPMP, the Multi-Compartment Segmentation plugin plays a crucial role in segmenting six major macro-anatomical structures in WSIs, generating AI-driven annotations that are immediately accessible to pathologists for visualization. After undergoing a structured review and quality control (QC) process, pathomic features are extracted from these annotations and uploaded to the KPMP Atlas, thereby enhancing accessibility for the broader research community.

Moreover, ComPRePS integrates human expertise into the AI workflow, forming a Human-AI loop that continuously improves as more users engage with the platform. Human supervision is central to the successful implementation of AI in medicine, as it ensures reliability, safety, and ethical application of technology. Human oversight functions as an essential safeguard for error correction in AI/ML systems, while preserving much of the efficiency these tools provide in clinical decision-making^54^. Supervision also builds trust among healthcare professionals, regulators, and patients by ensuring transparency and accountability^55,56^. Our Human–AI quality control loop is designed to be user-friendly, enabling pathologists to seamlessly review segmentation results as part of their workflow. This process not only corrects abnormal findings and reinforces confidence in the output classifications but also generates valuable feedback for model retraining, thereby supporting the continuous refinement of the computational pipelines.

ComPRePS is the only open-sourced platform that we are aware of exemplifying the close collaboration among algorithm developers, medical researchers, and clinicians that is necessary to bring the latest and greatest AI into bedside and patient care. Developers continuously iterate algorithms and integrate cutting-edge AI models - performing automation and solving tasks at scale. Researchers, in turn, bring rigor and analytical depth to evaluate and validate AI models across diverse kidney diseases - generating new scientific hypotheses and mechanistic insights of pathological impact. Clinicians and physicians apply expert domain knowledge and supervisory oversight to guide diagnosis, optimize treatment decisions, and ultimately define the trajectory of patient outcomes. By integrating the efforts of three expert groups that traditionally operate independently - AI developers, medical researchers and hospital clinicians - ComPRePS creates an evolutionary opportunity of taking histopathological evaluation from a subjective, experimental technology to quantifiable, reproducible clinical action-making workflow that could fulfill the prospect of precision medicine.

We have presented two case studies that collectively demonstrate the versatility and impact of ComPRePS in both clinical and research settings.

*In the first case*, ComPRePS significantly reduced interobserver variability and improved biopsy scoring accuracy across all lesion types, enabling pathologists, regardless of their expertise level, to achieve expert-level consistency through its AI-assisted human–computer interactive interface. This capability is particularly valuable for procurement biopsy assessments, where the limited availability of expert renal pathologists and variability among on-call evaluators can directly influence transplant decision-making. Recently published expert recommendations on the responsible use of AI in kidney care reinforce our view that clinician oversight must remain central to AI development and deployment. In our study, this principle is reflected in the ability of AI-assisted procurement biopsy interpretation to support more confident and precise donor kidney assessment, with the potential to expand organ utilization by avoiding unnecessary discards.^57^

The reproducibility of different histological lesions in donor biopsies in our study was like that reported by Azancot *et. al*.^58^ in their evaluation of renal biopsies from expanded criteria donors. For glomerulosclerosis, weighted κ was 0.76 in our cohort versus 0.86; for intimal thickening, κ was 0.47 versus 0.37. Since IFTA was scored as a single category in our study, we do not have a direct comparator; however, the reported values for interstitial fibrosis (κ = 0.31) and tubular atrophy (κ = 0.14) are close to our combined IFTA score (κ = 0.29). After incorporating a computational tool, reproducibility improved substantially: glomerulosclerosis increased by +0.10 κ, intimal thickening by +0.20 κ, and IFTA by +0.28 κ. The improvement was even more pronounced when comparing the average agreement between RP1 and RP2 with that of the on-call pathologists (OP), showing gains of +0.20 κ for glomerulosclerosis, +0.24 κ for intimal thickening, and +0.27 κ for IFTA. Ensuring reproducibility across existing classification systems remains a major challenge in Nephropathology. This challenge is not unique to procurement biopsy assessment. Popular scoring tools such as the Banff classification for allograft pathology, the MEST-C score for IgA nephropathy, and the ISN/RPS classification for lupus nephritis have long faced similar issues. While these systems offer good overall reproducibility when applied by experienced nephropathologists, their consistency tends to diminish in the hands of less experienced evaluators. This variability can impact diagnostic accuracy, therapeutic decisions, and ultimately patient outcomes, highlighting the need for tools that enhance consistency and support broader implementation across different levels of expertise. Beyond clinical applications, this case study also highlights the platform’s potential in research and education. The tools developed for Remuzzi score calculation, and the flexible workflows enabled by ComPRePS allow computational data scientists and clinical researchers to collaboratively design, refine, and implement AI-supported tools in a dynamic, iterative manner.

*In the second case*, ComPRePS facilitated high-throughput, quantitative morphological analysis of glomeruli and tubules across amyloid nephropathy, diabetic nephropathy, and minimal change disease. Automated multi-compartment segmentation and feature extraction enabled precise measurements - such as eosinophilic area fraction and texture metric-revealing disease-specific morphological patterns and statistically significant inter-group differences. These insights, which would be difficult to capture with manual, semi-quantitative methods, illustrate how ComPRePS bridges descriptive and predictive pathology, supporting reproducible, explainable, and biologically interpretable analyses.

Our case studies also highlight limitations that underscore the need for continued development of the platform. In the transplant setting, the widespread reliance on frozen sections for procurement biopsy assessment, driven by time constraints, poses a significant challenge. The performance and generalizability of our tools must be validated in this context, where tissue quality and staining consistency may differ from formalin-fixed samples. In native kidney diseases, multiple stains are often required to accurately identify specific structures or pathological components, such as the tunica media or amyloid deposits, and our current approach may be limited without integration of multimodal staining. In addition, the native disease cohorts analyzed here involve morphologically less complex entities compared with conditions such as IgA nephropathy or lupus nephritis. These disorders exhibit a broader spectrum of overlapping and spatially heterogeneous lesions, which may pose additional challenges for automated segmentation and feature extraction. Future work must therefore evaluate ComPRePS in disease contexts with higher morphological complexity to ensure robust generalizability. Finally, while our work has primarily focused on chronic injury and glomerular diseases, future development should also address acute conditions, such as acute interstitial nephritis, rejection in kidney transplantation, acute tubular necrosis, and other forms of acute kidney injury, where prompt and accurate histopathological interpretation is crucial for effective clinical decision-making. Restricted cohort size and limited pathologist diversity are also among limitations of this study; expanding the donor demographics and expertise range of reviewing pathologists will be critical to ensuring the consistency and generalizability of our findings.

Together, these examples underscore ComPRePS’ potential to standardize lesion assessment, accelerate morphological research, and enhance decision-making in both routine clinical practice and advanced Nephropathology studies, while also pointing to clear opportunities for expansion and refinement.

As future work, we aim to expand the platform in the following ways:

1. Empowering researchers with AI-assisted tools and scalable computing infrastructure to accelerate scientific discovery.
2. Facilitating intuitive learning for students, encouraging curiosity and rapid skill development.
3. Deploying ComPRePS in real-world clinical settings to support pathologists in diagnostic and research workflows.
4. Expanding the model zoo to cover additional renal structures and associated histopathological pathomics providing a platform for broader understanding of complete kidney health and clinical holistic treatment planning.
5. Strengthening interpretability and medical explainability for clinicians and doctors by ensuring AI-derived insights are transparent, actionable, and can be summarized to patients.
6. Integrating multi-modal data – molecular, genomic, immunohistochemistry/immunofluorescence and granular electronic health records (EHR) for determining current patient condition and accurate predictive modeling of disease prognostics.
7. Continuously establishing ComPRePS as a global hub, developing tool features, new visualization routines, reducing pain-points for developers to bring their AI models into production, and empowering researchers/clinicians to apply and validate AI into novel use-cases.

By continuously refining ComPRePS and fostering a growing user community, we aim to make AI an indispensable asset in both research and clinical practice.

## Methods

### Ethics statement

This study complied with all applicable ethical guidelines and was approved by the Institutional Review Boards of the University of Florida (UF; IRB202300413) and the University of Coimbra (UC; IRB00003904). All human tissue samples were collected under IRB-approved protocols from participating institutions. Samples were fully de-identified before being accessed or analyzed. Informed consent was obtained by contributing institutions for the native kidney samples, and participants did not receive any form of compensation.

### ComPRePS: A next-generation collaborative ecosystem for Computational Pathology

ComPRePS is a cloud-enabled, horizontally scalable platform designed to unite clinicians, researchers, and AI model developers around the shared goal of extracting robust, actionable insights from WSIs. Our previous system, HistoCloud^35^, established a solid foundation and demonstrated success in research environments, but large-scale real-world deployment in the clinical workflow had not been tested, due to bottlenecks in terms of functionality, computational throughput, scalability, and PHI data security. Beyond improvements in performance and scalability, we have substantially expanded ComPRePS with new analytical capabilities and user-facing tools. The platform now features an extensive model zoo, incorporating newly developed AI models for kidney diseases, including arterial intima and peritubular capillary (PTC) segmentation, along with an overhauled suite of visualization utilities. It also introduces enhanced modules for granular feature extraction, pathomic quantification, and disease classification.

As collaborations expanded, the need for diverse AI models, advanced analytics, and scalable computation increased. New pipelines for segmentation, feature extraction, and disease classification enabled richer analyses but also imposed heavier computational demands for large-scale WSI processing. To meet these demands, we needed a platform capable of running multiple AI workloads efficiently and concurrently across hundreds of slides. The original HistoCloud platform relied on monolithic architecture and was limited to deployment on a single server or VM (virtual machine), where performance was constrained by the available computation and memory of that machine. Scaling up resources such as Central Processing Units (CPUs), Graphical Processing Units (GPUs), or Random Access Memory (RAM) required vertical scaling, i.e., adding more capacity to the same system, quickly became a bottleneck and restricted responsiveness to user needs. ComPRePS extends this to a massively parallel, distributed computing framework across a HPC environment, enabling efficient data crunching and high-throughput AI/ML analysis across hundreds of WSIs concurrently.

To address the limitations of HistoCloud and to build a truly next-generation tool for computational pathology, we re-engineered the entire platform architecture to create ComPRePS, the first tool of its kind to the best of our knowledge. Our focus was to deliver a robust, powerful, and future-proof system that puts users at the center of the experience.

Figure 7 presents the redesigned ComPRePS system architecture. The new system uses microservice-based designs, in which modular building blocks can evolve independently. This means that each component can be updated, scaled, or replaced without disrupting the rest of the system, providing flexibility as user needs, and collaborations continue to grow.

**Figure 7.**
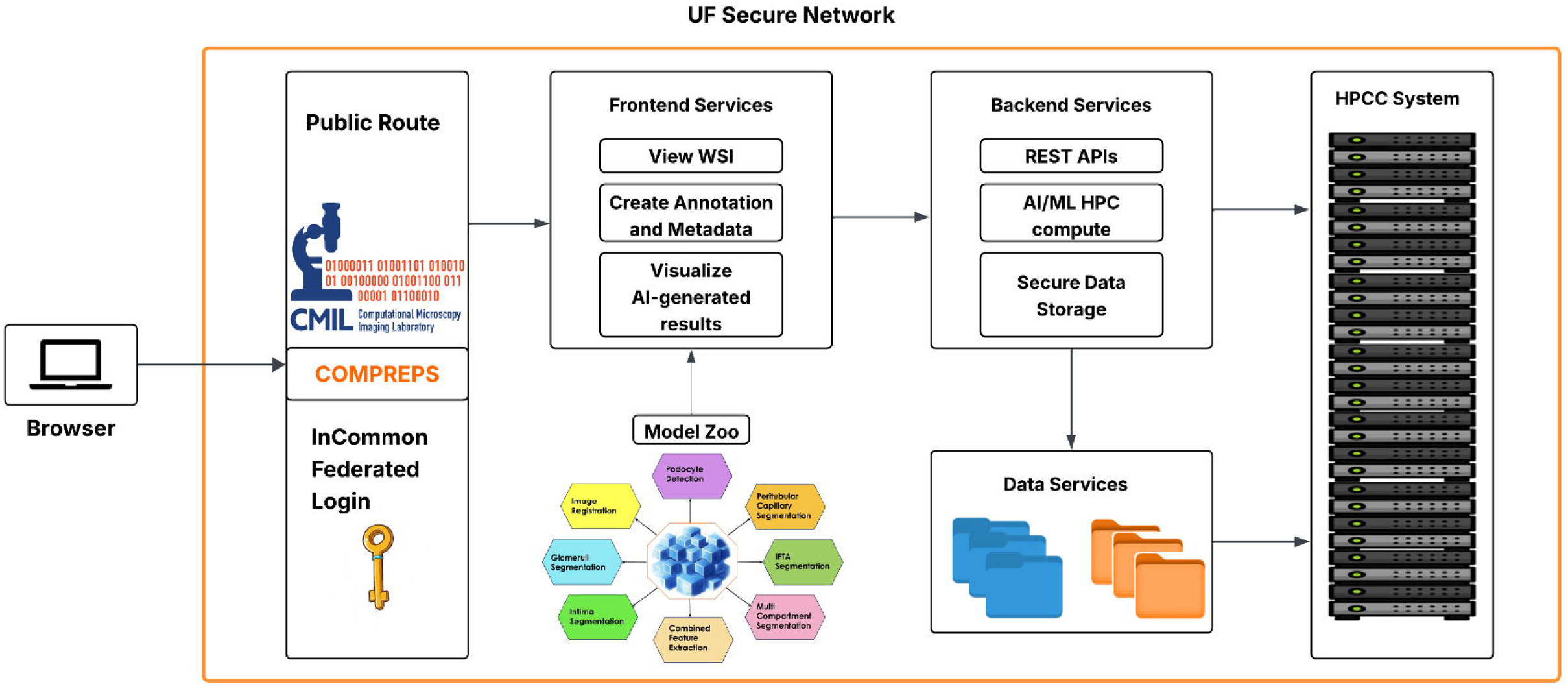
Architecture diagram of ComPRePS illustrating the major building blocks. From left to right, the diagram shows the Authentication layer, various services (frontend, backend, and data services), and the Model Zoo. The integration with the HiperGator Supercomputer, UF’s high-performance computing system, is also depicted. Abbreviations: AI/ML, artificial intelligence/machine learning; API, application programming interface; CMIL, Computational Microscopy Imaging Laboratory; HPC/HPCC, high-performance computing (center); REST, representational state transfer; UF, University of Florida; WSI, whole-slide image.

From a user’s perspective, the workflow begins with data ingestion, where WSIs, structured clinical data, and pathologists’ diagnostic notes including reported structural injuries and abnormalities can be seamlessly uploaded via ComPRePS user-interface. Once the data is in the system, users can start to visualize WSIs and results, create or evaluate annotations for desired quality, and more. Users can then launch AI pipelines from the model zoo for tasks like segmentation. The results are then displayed to the user and ingested into the database for persistent storage as well as can be added to the user’s original location allowing downstream analysis.

Below we describe the individual services that ComPRePS offers to support this end-to-end workflow.

### Front-end Services

The front-end services of ComPRePS were designed to provide a user-friendly interface while retaining analytical depth. The UI played a significant role in the success of HistoCloud, so we have retained its key strengths while introducing targeted enhancements. Some of the new enhancements include supporting more image formats to aid and keep up with the evolving landscape of imaging technologies, and the plotting tool has been overhauled. The plotting tool now supports importing clinical metadata from multiple sources - including slide-level data, AI analysis results, folder-level data, cohort-level data, and annotation-level data. In addition, it offers new visualization options such as violin plots and line graphs, along with several usability improvements like a search bar, detailed filters, free-form selection, and other enhancements. More information regarding the front-end services is described in the *Supplementary Information*.

### Back-end Services

The backend services form the computational core of ComPRePS, which is now distributed and able to leverage HPC resources^36^. In practical terms, HPC refers to the ability to connect many powerful computers so that complex tasks, such as running AI models on hundreds of WSIs, can be processed in parallel. In our earlier version, HistoCloud, all processing was confined to a single server, which limited performance and flexibility. In ComPRePS, each analysis request automatically receives dedicated computing resources, ensuring multiple users can run demanding AI workloads simultaneously without delays. Once a task is complete, those resources are released for others to use, allowing the system to operate efficiently and maintain consistent performance.

Currently, ComPRePS is integrated with UF’s HiperGator HPC cluster, but it should also be compatible with any other SLURM-based HPC system (yet to be tested). This ensures broad portability for other institutions that wish to adopt the platform in case they have institutional restrictions on sharing data.

A major advancement in ComPRePS is its open and extensible design, which allows for seamless integration with other applications. In clinical and research environments, interoperability is essential, as multiple specialized tools need to work together to support end-to-end analysis. To enable this, ComPRePS provides standardized interfaces that allow external platforms to programmatically access its computing power and data management capabilities. These interfaces, known as application programming interfaces (APIs), serve as bridges that let different software systems communicate and exchange data automatically, without manual intervention. This capability is implemented through a set of Representational State Transfer (RESTful) APIs, which make it possible for external applications to submit AI tasks, retrieve results, or manage data directly through the platform. While This functionality was limited in HistoCloud, it is now actively used by external platforms such as FUSION^31^ and Kidease^37^, facilitating multi-faceted research and clinical workflows.

### Data Services

WSIs are often hundreds of megabytes to tens of gigabytes in size, making data management a central challenge. ComPRePS integrates directly with UF’s HiperGator supercomputing environment, which employs a Lustre-based parallel network file system^38^, a standard architecture in many HPC deployments. This setup provides multiple storage tiers optimized for speed, capacity and long-term data retention, allowing for efficient handling of both active and archived datasets. Through symbolic linking, users can connect existing data directories directly to the platform, avoiding data duplication and enabling flexible pre- and post-processing tasks.

Collaboration across institutions is supported by Globus^39^, which enables secure and high-speed transfers of large datasets. ComPRePS can symbolically link transferred data into its workflows immediately after upload, allowing collaborators to begin analysis without delay. Metadata management has been built into the data layer, permitting users to attach metadata at the slide, dataset, or algorithm level and import external annotations for downstream analyses.

### Security Services

Security and authentication are fundamental aspects of ComPRePS. The platform now integrates with the InCommon Federation, a trusted identity gateway for the US research and education community. This provides secure single sign-on (SSO) for both local and cloud-based services. Additionally, two-factor authentication and virtual private network (VPN) locking ensure the security of both HPC resources and PHI data^40^. Further details on these security measures are provided in the *Supplementary Information*.

### Model Zoo Ecosystem

What makes ComPRePS so powerful is the creation of the collaborative Model Zoo, which has become a hub for sharing and deploying AI pipelines. The earlier HistoCloud system contained a few internally developed models, such as glomeruli and IFTA segmentation. However, it did not provide a scalable framework ingesting a growing range of AI pipelines from external contributors. ComPRePS now hosts a growing set of AI pipelines contributed by external groups, such as PTC segmentation by Duke and Emory University^17^, and cell segmentation by DeepCell^18^, reflecting the explosive contribution of AI from the global community and maturing into an “all-in-one” ecosystem. Summaries of key plugins are provided below, while expanded descriptions are available in the *Supplementary Information*.

### Multi-Compartment Segmentation

This pipeline integrates a panoptic segmentation model to segment 6 kidney FTUs, which includes non-globally sclerotic glomeruli, globally sclerotic glomeruli, tubules, cortical and medullary interstitium, and arteries and arterioles^41^. FTU segmentation masks are stored in a lightweight JSON format for easier interoperability to downstream applications and displayed as layers within the UI, enabling large-scale reproducible FTU-level segmentation across datasets.

### Combined-Feature Extraction

The combined-feature extraction plugin automates the extraction of various categories of morphometric and pathomic features derived from each segmented structure^42^. Our team brings multi-disciplinary expertise in image pathomics - a data-driven methodology for systematically extracting vast quantitative information from kidney tissues. The comprehensive library has features ranging from ready-made clinically interpretable to subtle tonon-obvious patterns that are beyond the perceptual capabilities of the human visual system has been instrumental in elucidating biomarker discovery and understanding diagnostic outcome. In ComPRePS, we have implemented highly efficient computer vision (CV) pipelines spanning multiple levels of complexity, from clinically interpretable morphological and color-based descriptors (e.g., compartment dimensions, nuclear shape, and stain distribution) to advanced textural and spatial metrics that characterize tissue heterogeneity and spatial relationships among sub-compartments such as eosinophilic regions, lumens, and nuclei within the renal microenvironment. Results can be visualized as scatter or violin plots within the UI, supporting both single-slide inspection and cross-slide comparisons and stored for further downstream analysis.

### IFTA Segmentation

The IFTA segmentation plugin implements a convolutional neural network (CNN) model trained on multi-institutional kidney biopsies^16^. The tool provides reproducible, quantitative assessments of cortical fibrosis and tubular atrophy that correlate with patient outcomes, reducing reliance on subjective visual scoring.

### PTC Segmentation

The PTC segmentation plugin integrates an externally developed model to identify and quantify peritubular capillaries, which play a critical role in chronic kidney disease (CKD) progression. Duke and Emory University researchers have built a well validated deep learning (DL) network for cortical PTC segmentation using 280 PAS-stained biopsies from the Nephrotic Syndrome Study Network^17^. This model has been incorporated into ComPRePS as a ready-to-use plugin, expanding the accessibility and outreach of plug-and-play AI tools by enabling researchers worldwide to apply validated models easily to their own datasets. The plugin highlights the interoperability of ComPRePS by enabling the reuse of validated external pipelines in a standardized form and opens the potential of incorporating more accurate models for any task as AI evolves at an unprecedented pace.

### Podocyte Detection

Renal diseases often result in podocyte injury, making quantification of podocyte loss clinically significant for assessing and tracking disease progression. Currently, podocyte detection relies on manual identification from PAS-stained renal sections - a process that is highly subjective, time-consuming and exhibits high interobserver variability. To overcome these limitations, we developed and integrated multiple AI-enabled pipelines within ComPRePS for automated, objective, and scalable podocyte detection and quantification, enabling consistent assessment across patient cohorts. The PodoCount plugin^43^ detects podocyte nuclei from renal immunohistochemistry images by applying specific podocyte-specific antibodies. We have also integrated the CNN based PodoSighter plugin^43,44^ to identify podocytes from routine brightfield renal images using the standard PAS counterstained with hematoxylin. The plugins support direct quantification through ComPRePS UI, enabling large-scale podocyte analysis without requiring local computational infrastructure and enhancing the possibility of advanced causation-correlation analysis of podocyte displacement and loss in relation to glomerular disease progression.

### Image Registration

The registration plugin aligns WSIs across modalities (for example, histology and spatial molecular imaging) or serial sections using rigid, affine, or thin plate spline transformations. Landmark-based and intensity-based options are provided, with outputs automatically stored for downstream integration. This enables robust cross-modality comparisons and correction of common histological artifacts.

### Arterial Intimal Fibrosis Computation

Thickening of arterial intima – the innermost layer of the arterial wall – accompanied with narrowing of the arterial lumen provides a quantitative marker of vascular injury, sclerosis and fibrosis deposition that are key features of progression of CKD. Accurate, automated quantification of intimal thickness enables precise and objective assessment of severity of vascular pathology. We have developed and integrated an attention U-Net pipeline, introducing learnable attention mechanism into the traditional encoder-decoder style network to prioritize clinically meaningful lesion segmentations (manuscript in preparation) to compute arterial intimal thickness, lumen reduction, and stenosis. Arteries are cropped, processed, and analyzed, with results presented as both numerical metrics and visual overlays. The pipeline facilitates evaluation of vascular remodeling at a large scale within kidney biopsies, allowing for correlation with clinical markers and long-term renal function decline.

### Interoperability: Glomeruli Segmentation Models

In addition to our in-house multi-compartment model, ComPRePS hosts several externally developed glomerular segmentation pipelines incorporating state-of-the-art AI architectures, including a U-Net–based model for frozen tissue sections^45^ and a SegFormer-based model trained on The Human Biomolecular Atlas Program (HuBMAP) and Human Protein Atlas (HPA) datasets^19,46^. Both open-sourced pipelines^47^ are containerized as ComPRePS plugins and made ready-to-use within the platform, demonstrating the ability to integrate community-developed tools with minimal modification and quickly bring latest and greatest AI through ComPRePS to clinicians.

### Classification

ComPRePS includes a supervised classification pipeline that predicts clinical outcomes from histopathological WSIs. This approach uses morphometric and pathomic features extracted through the ComPRePS segmentation pipeline as inputs to an optimized XGBoost classifier^48–50^, enabling an efficient and interpretable end-to-end workflow, from tissue segmentation and feature extraction to outcome prediction. Showcasing its broader applicability, we evaluated ComPRePS for predicting treatment response in lupus nephritis using morphometric features derived from segmented glomeruli. While these analyses are exploratory and not yet published, the initial results are promising, demonstrating the platform’s translational potential to support kidney precision medicine approaches across diverse chronic kidney diseases. We aim for a dedicated publication detailing this application and expanding our findings.

### HistoQC

HistoQC^51^ is an externally developed, open-source pipeline for automated quality control of WSIs. It provides a suite of modules to assess image quality factors - artifact detection, staining variability, contrast, brightness, blurriness, and the presence of slide-preparation artifacts like folds or pen markings – ensuring consistency and reproducibility in large-scale histopathological image analysis. Within ComPRePS, we have integrated HistoQC as part of the image pre-processing module prior to compartmental segmentation and feature extraction, ensuring computational quality assurance and robustness of downstream analyses.

### Clinical Use - Case 1: Assessment of Donor Biopsies for Kidney Transplantation Using ComPRePS

#### Study Data and Design

We analyzed a cohort of 213 whole-slide images (WSIs) from procurement core needle biopsies of 152 deceased kidney donors, collected between 2011 and 2023 at the Kidney Transplant Unit of the Coimbra Hospital and University Center in Portugal. Detailed donor demographic and clinical variables are presented in Supplementary Table 14.

The dataset included both transplanted kidneys (*n* = 139) and discarded kidneys (*n* = 74), as determined by on-call pathologists using the Remuzzi score, to capture a broad spectrum of pre-existing chronic lesions. We excluded biopsies lacking acceptable staining quality, autopsy artifacts, or representation of at least 10 glomeruli and 2 arteries, as determined by on-call pathologists. Tissue sections (2 μm thick) were prepared via a 3-hour fast Formalin-Fixed Paraffin-Embedded (Fast-FFPE) method, a rapid protocol to accelerate tissue processing and preserving morphological integrity of kidney structures and stained with PAS (detailed description provided in Supplementary Information). All slides were digitized at 40X magnification (0.25 μm resolution /pixel) using the Aperio CS2 scanner (Leica Biosystems, Vista, CA, USA).

#### Biopsy Evaluation Workflows

Procurement biopsies were evaluated using a modified Remuzzi’s classification, the histopathological grading system routinely adopted by the University of Coimbra Transplant Center for donor kidney assessment ^23^. Instead of the original Remuzzi method for estimating arterial and arteriolar narrowing as a measure of vascular chronicity, pathologists applied the internationally standardized Banff Lesion Score cv visual analog scale, as defined by the Banff Classification ^25^. Traditional Remuzzi arterial scores were not assigned or used in subsequent analyses.

We compared four distinct biopsy evaluation workflows:

- Original on-call pathologist (on-call: OP): Standard clinical assessment.
- Reviewer pathologist (Reviewer: RP): Independent evaluation by 2 experienced nephropathologists.
- Computational method (ComPRePS): Fully automated end-to-end AI-driven analysis by ComPRePS.
- Human–AI collaboration, designated as the ground-truth (GT) comparator (Reviewer-ComPRePS: GT): Experienced pathologists review augmented by ComPRePS’s kidney compartment segmentations.

A detailed description of the Remuzzi score calculation by on-call and reviewer pathologists is provided in the *Supplementary Information*.

The computational Remuzzi score (ComPRePS) was obtained through the following steps:

i. Global glomerulosclerosis percentage – We applied the multi-compartment segmentation plugin to classify two glomerular types within each WSI: non-globally sclerotic glomeruli and globally sclerotic glomeruli. The tool then quantified the number of glomeruli in each class and computed the glomerulosclerosis fraction.
ii. Interstitial fibrosis and tubular atrophy (IFTA) – We applied the IFTA estimation plugin in ComPRePS to segment IFTA regions, defined as patchy areas of interstitial fibrosis and tubules exhibiting atrophic changes. These segmented regions were quantified based on the fraction of the cortical IFTA area relative to the total cortical area. The method for calculating the cortical area of each WSI and for converting computational IFTA measurements into Remuzzi interstitial fibrosis (IF) and tubular atrophy (TA) scores is provided in detail in the *Supplementary Information*.
iii. Arterial intimal fibrosis – The computational pipeline cropped for each selected artery and applied the intima segmentation model to generate a predicted intimal mask. Given the irregular and distorted morphology of the intima in most cases, our objective was to calculate radial thickness across the segmented mask, defined as the distance from each pixel on the inner intimal boundary to its nearest pixel on the outer boundary. The mean intimal thickness was then derived as the average of all radial thicknesses. In addition, we quantified fibrointimal thickening using the stenosis ratio, defined as the ratio of the intimal area to the sum of the intimal and luminal areas. To compute these areas, we employed a computer vision strategy based on contour detection applied to the predicted intimal mask.
iv. Finally, the individual lesion values for glomerulosclerosis, IFTA, and intimal fibrosis were converted to lesion scores using predefined thresholds (Supplementary Table 15) and combined to compute the computational Remuzzi grade.

For the Human–AI collaboration (Reviewer–ComPRePS) Remuzzi score calculation, the Reviewer assessed each lesion category by visualizing ComPRePS segmentations and individual score predictions through ComPRePS UI. The Reviewer would oversee performing AI quality assurance and could refine predictions as per liking to improve accuracy. For glomeruli, this included adding or removing false-positive and false-negative annotations for non-sclerotic and globally sclerotic glomeruli. For IFTA, the Reviewer could confirm or adjust IFTA boundaries when necessary or choose to delete the lesion annotations altogether. For arterial intimal fibrosis, the Reviewer selected the artery or arteries considered most representative of severe intimal thickening for morphologic analysis. ComPRePS generated fibro-intima predictions and stenosis ratios on chosen arteries across multiple arterial orientations and varying degrees of thickening. The Reviewer could then select the region of intimal thickening that best corresponds to the tissue section or choose to accept the ComPRePS-generated mean intimal thickness. This is particularly valuable in arterial longitudinal sections, where using a mean stenosis ratio alone can be misleading.

### Clinical Use - Case 2: Quantitative Morphological Analysis in a Kidney Disease Cohort

#### Image data and Feature Processing

The dataset consisted of WSIs of kidney biopsy samples for three different CKD groups - Amyloid Nephropathy (AN), Diabetic Nephropathy (DN), and Minimal Change Disease (MCD). Our cohort comprised a diverse, multi-institutional set of cases from the University of Florida (Gainesville, FL), Johns Hopkins University (Baltimore, MD), and the University of Michigan (Ann Arbor, MI). Data collection was conducted following research protocols approved by the Institutional Review Boards of the participating institutions. Each disease group included six WSIs, which were used for inter-group analysis. Dataset preprocessing was first performed using HistoQC. Glomeruli and tubules were segmented using the multi-compartment segmentation plugin for each WSI. Although multi-compartment segmentation isolates additional kidney structures, this study focused solely on glomerular and tubular annotations for simplicity. Morphological and pathomic features were subsequently extracted from these structures using the feature extraction pipeline ^53^ available in ComPRePS. Detailed descriptions of the pathomics extracted for this use-case are provided in the *Supplementary Information*. Following feature extraction, feature visualization was performed between disease groups for statistical comparison and cluster analysis. This plugin outputs two files: (i) a derived metadata file for plotting and metadata mapping in the Metadata Plot tool, and (ii) an annotation file that enables direct visualization of the identified pathomic clusters on the WSIs. Full details of the Feature Visualization plugin and associated file formats are provided in the *Supplementary Information*.

### Statistics and Reproducibility

#### Clinical Use - Case 1: Assessment of Interobserver Agreement of the Remuzzi Scoring

To assess performance and agreement between the different observers, we analyzed each Remuzzi lesion score as well as the overall score. The evaluation comprised: (i) confusion matrices comparing ComPRePS predictions with the Reviewer-ComPRePS pipeline (GT), as well as comparisons of GT with on-call assessments; (ii) prediction metrics – including precision, recall, F1 score, and accuracy - of ComPRePS relative to on-call pathologists, using the GT as the reference standard; and (iii) interobserver agreement, quantified using Cohen’s Kappa for pairwise comparisons between pathologists and ComPRePS biopsy evaluation workflows across all Remuzzi lesions. Cohen’s Kappa was selected as it accounts for chance agreement, providing a more robust and reliable measure than simple percent of agreement.

#### Clinical Use- Case 2

Summary statistics were reported as mean, median, and interquartile range. Feature distributions were assessed visually using histograms and violin plots and tested for normality using a chi-square goodness-of-fit test. Associations between glomerular eosinophilic area fraction and total glomerular area were evaluated using linear regression, with strength of association quantified by the coefficient of determination (R²). Between-group differences were assessed using one-way ANOVA with Tukey’s honestly significant difference (HSD) post-hoc correction for multiple comparisons when appropriate. All statistical visualizations were generated using ComPRePS’s integrated visualization tools. Statistical significance was defined as a two-sided *p* < 0.05.

## Data Availability

All WSIs, generated metadata, and AI segmented renal compartments supporting the clinical use-case 1 and 2 are available for interactive viewing through the ComPRePS Computational Microscopy Imaging Lab Slide Archive at https://athena.rc.ufl.edu (Collections/Journals/ WSI_PaulRodrigues_NatureComm_2026). Detailed guidance for navigating WSIs and computational annotations is provided in the Supplementary Information in section “*Navigating WSIs with Annotation Visualizations through ComPRePS UI*”.

## Code Availability

All software components developed and utilized in this study are publicly accessible through the ComPRePS ^52^ Wiki (https://compreps-wiki.cmilab.nephrology.medicine.ufl.edu/), which provides detailed documentation and usage examples. The repository includes the high-performance computing (HPC) integration pipelines that support large-scale segmentation, feature extraction, and model training on the HiperGator cluster.

A supplementary video in the Wiki demonstrates the main functionalities of ComPRePS, including (a) loading and analyzing WSIs through the web interface, (b) retrieving data from the HiperGator server across the frontend and backend environments, (c) executing segmentation pipelines for all tissue compartments, (d) performing feature extraction, and (e) mapping selected features for visualization and classification.

## Acknowledgement

The authors would like to acknowledge the support of the following grants, which made this work possible: R01 DK114485, R21 DK128668, R01 DK129541, OT2 OD033753, OT2 OD038014 and U01 DK133090.

We also extend our sincere thanks to Samantha Hoffman, Christina Beharry, and Audrey Singletary for their valuable assistance in reviewing and editing the manuscript to ensure its clarity and quality and for Dr. Laura Barisoni for providing us with the PTC Segmentation Model.

## Multi-Institutional and Interdisciplinary Collaboration

This study was conducted through a close, sustained collaboration between the University of Florida (UF) and the University of Coimbra (UC), integrating complementary expertise across computational systems engineering, artificial intelligence, data science, nephropathology, and clinical transplantation. System developers and data scientists from UF led ComPRePS platform architecture and AI implementation, nephropathologists from UC provided domain-driven validation and clinicians from UC participated in human-AI workflow and guided clinical translational applicability.

## Author Contributions

A.S.P. led the design, deployment, and validation of the ComPRePS computational pathology platform, including AI pipeline integration, scalable infrastructure, resource management, and overall technical supervision beside drafting and editing the manuscript. L.A.R. led clinical conceptualization and evaluation, including study design, methodology, cohort curation, clinical validation, and manuscript drafting and editing. S.K.C.K., S.D., H.A., and O.M. contributed to scalable system architecture, algorithm implementation, AI model optimization, and deployment to UF infrastructure. D.M., W.D., and S.P.B. supported infrastructure and feature development, usability-focused engineering, and implementation of data visualization approaches. C.Pa., C.Pi., V.S., A.F., and R.A. contributed to annotation generation, AI model validation, human-AI workflow, quality control and biological interpretation refinement. A.Z.R., A.F. and B.A.S. provided expert nephropathology oversight and contributed to manuscript review. P.S. conceptualized the idea of ComPRePS, supervised the whole study, provided senior leadership for the computational and translational framework, mentored the team, and critically revised the manuscript.

## Competing Interests

P.S. and A.S.P. serve on the advisory board of DigPath Inc. Cloud-CIPHERS - Cloud services for Computational AI Infused Pathology harnessing Histologic platform which encompasses ComPRePS is protected by ©Copyright 2024 University of Florida Research Foundation, Inc. All Rights Reserved. *FUSION* software is protected by ©Copyright 2023-25 University of Florida Research Foundation, Inc. All Rights Reserved. The remaining authors declare no competing interests.

## Regulatory Information

This study complies with all relevant ethical regulations and was approved by the Institutional Review Boards at the University of Florida and at the University of Coimbra.

## Notes

https://compreps-wiki.cmilab.nephrology.medicine.ufl.edu/)

### Summary of Updates

In this revision, the author information was updated, based on the authorship contributions in restructuring and rewriting the manuscript knowledge as the platform progressed in development.

