## Supplemental Information for "*ComPRePS*: Unlocking Scalable AI Analysis for Computational Renal Pathology"

#### *ComPRePS*: Unlocking Scalable AI Analysis for Computational Pathology in Clinical Applications

#### Clinical Use-case 1 Results: Comprehensive Analysis of Interobserver Agreement for Each Remuzzi Score Component

*Supplementary Table 1:* Confusion matrix of glomerulosclerosis grade (0-3): ComPRePS and On-call against GT

(a) Confusion Matrix comparing glomerulosclerosis ground truth (GT) scores with ComPRePS predictions. Shaded cells indicate the percentage of correct predictions

| | | ComPRePS | | | |
|---|---|---|---|---|---|
| | | **0** | **1** | **2** | **3** |
| **GT** | **0** | 94.7% | 2.3% | - | - |
| | **1** | 3.5% | 97.0% | 18.2% | - |
| | **2** | 1.8% | 0.8% | 77.3% | - |
| | **3** | - | - | 4.5% | 100.0% |

(b) Confusion Matrix comparing glomerulosclerosis ground truth (GT) scores with On-call pathologist's assessments. Shaded cells indicate the percentage of correct predictions

| | | On-call | | | |
|---|---|---|---|---|---|
| | | **0** | **1** | **2** | **3** |
| **GT** | **0** | 94.7% | 17.2% | 3.5% | - |
| | **1** | 5.3% | 79.9% | 42.1% | - |
| | **2** | - | 3.0% | 52.6% | 33.3% |
| | **3** | - | - | 5% | 66.7% |

*Supplementary Table 2:* Performance metrics between ComPRePS and On-call for glomerulosclerosis assessments

(a) ComPRePS against GT

| | **Precision** | **Recall** | **F1** | **Accuracy** |
|---|---|---|---|---|
| **0** | 0.95 | 0.95 | 0.95 | 0.97 |
| **1** | 0.97 | 0.96 | 0.96 | 0.95 |
| **2** | 0.77 | 0.90 | 0.83 | 0.97 |
| **3** | 1.00 | 0.67 | 0.80 | 0.99 |

(b) On-call against GT

| | Precision | Recall | F1 | Accuracy |
|---|---|---|---|---|
| **0** | 0.69 | 0.95 | 0.80 | 0.87 |
| **1** | 0.91 | 0.80 | 0.85 | 0.82 |
| **2** | 0.67 | 0.53 | 0.59 | 0.93 |
| **3** | 1.00 | 0.67 | 0.80 | 0.99 |

*Supplementary Table 3:* Confusion matrix of the IFTA grade (0-6): ComPRePS and On-call against GT

(a) Confusion Matrix comparing IFTA ground truth (GT) scores with ComPRePS predictions. Shaded cells indicate the percentage of correct predictions

| | | **ComPRePS** | | |
|---|---|---|---|---|
| | | **0** | **2** | **4** |
| **GT** | **0** | 75.0% | 7.3% | - |
| | **2** | 25.0% | 90.7% | - |
| | **4** | - | 2.0% | - |

(b) Confusion Matrix comparing IFTA ground truth (GT) scores with On-call pathologist's assessments. Shaded cells indicate the percentage of correct predictions

| | | **On-call** | | |
|---|---|---|---|---|
| | | **0** | **2** | **4** |
| **GT** | **0** | 34.6% | 12.6% | - |
| | **2** | 65.4% | 81.4% | 25.0% |
| | **4** | - | 6.0% | 75.0% |

*Supplementary Table 4:* Performance metrics between ComPRePS and On-call for IFTA assessments

(a) ComPRePS against GT

| | **Precision** | **Recall** | **F1** | **Accuracy** |
|---|---|---|---|---|
| **0** | 0.75 | 0.91 | 0.82 | 0.75 |
| **2** | 0.91 | 0.78 | 0.84 | 0.91 |
| **4** | - | 0.00 | - | 0.98 |

(b) On-call against GT

| | **Precision** | **Recall** | **F1** | **Accuracy** |
|---|---|---|---|---|
| **0** | 0.35 | 0.73 | 0.47 | 0.35 |
| **2** | 0.81 | 0.47 | 0.60 | 0.81 |
| **4** | 0.75 | 0.93 | 0.83 | 0.75 |

*Supplementary Table 5:* Confusion Matrix comparing intimal fibrosis ground truth (GT) scores with ComPRePS and On-call pathologist's predictions on the pathologist determined worst artery. For ComPRePS analysis, pathologists provided a particular angle for their selection of the worst artery along with guidance on where intimal thickness should be measured. ComPRePS computes intimal thickness along that direction to quantify intimal area and stenosis grade of the artery. Shaded cells indicate the percentage of correct predictions

(a) Confusion matrix: ComPRePS against GT. Shaded cells indicate the percentage of correct predictions

| | | **ComPRePS** | | | |
|---|---|---|---|---|---|
| | | **0** | **1** | **2** | **3** |
| | **0** | 85.7% | 7.1% | - | - |
| **GT** | **1** | 14.3% | 91.4% | 4.8% | - |
| | **2** | - | 1.4% | 93.7% | 16.7% |
| | **3** | - | - | 1.6% | 83.3% |

(b) Confusion matrix: On-call pathologist's assessments against GT. Shaded cells indicate the percentage of correct predictions

| | | On-call | | | |
|---|---|---|---|---|---|
| | | **0** | **1** | **2** | **3** |
| **GT** | **0** | 100.0% | 6.0% | 1.2% | - |
| | **1** | - | 81.0% | 55.3% | 15.2% |
| | **2** | - | 9.5% | 35.3% | 51.5% |
| | **3** | - | 3.6% | 8.2% | 33.3% |

*Supplementary Table 6:* Performance metrics between ComPRePS and On-call compared against GT for assessment of fibro-intima thickening

(a) ComPRePS against GT

| | **Precision** | **Recall** | **F1** | **Accuracy** |
|---|---|---|---|---|
| **0** | 0.86 | 0.55 | 0.67 | 0.96 |
| **1** | 0.91 | 0.94 | 0.93 | 0.94 |
| **2** | 0.94 | 0.91 | 0.92 | 0.94 |
| **3** | 0.83 | 0.96 | 0.89 | 0.96 |

(b) On-call against GT

| | **Precision** | **Recall** | **F1** | **Accuracy** |
|---|---|---|---|---|
| **0** | 0.65 | 1.00 | 0.79 | 0.95 |
| **1** | 0.57 | 0.81 | 0.67 | 0.64 |
| **2** | 0.55 | 0.35 | 0.43 | 0.60 |
| **3** | 0.52 | 0.33 | 0.41 | 0.79 |

*Supplementary Table 7:* Performance metrics between ComPRePS and On-call compared against GT for final Remuzzi grading of allocation/rejection decision for the organ transplant. The score threshold used by the Transplant Center for the allocation/rejection decision is as follows: grade 0-4: Allocate, grade 5: Physician/surgeon consultation, grade 6-12: Dual kidney transplantation/Discard

(a) ComPRePS against GT

| | **Precision** | **Recall** | **F1** | **Accuracy** |
|---|---|---|---|---|
| **0** | 0.50 | 0.33 | 0.40 | 0.98 |
| **1** | 0.75 | 0.86 | 0.80 | 0.93 |
| **2** | 0.17 | 0.22 | 0.19 | 0.89 |
| **3** | 0.62 | 0.62 | 0.62 | 0.84 |
| **4** | 0.66 | 0.73 | 0.69 | 0.77 |
| **5** | 0.81 | 0.63 | 0.71 | 0.80 |
| **6** | 0.55 | 0.63 | 0.59 | 0.89 |
| **7** | 0.50 | 0.86 | 0.63 | 0.95 |
| **8** | 0.67 | 0.67 | 0.67 | 0.99 |

(b) On-call against GT

| | Precision | Recall | F1 | Accuracy |
|---|---|---|---|---|
| **0** | 0.50 | 0.33 | 0.40 | 0.96 |
| **1** | 0.30 | 0.43 | 0.35 | 0.79 |
| **2** | 0.29 | 0.67 | 0.40 | 0.81 |
| **3** | 0.37 | 0.53 | 0.43 | 0.62 |
| **4** | 0.34 | 0.36 | 0.35 | 0.49 |
| **5** | 0.47 | 0.24 | 0.32 | 0.53 |
| **6** | 0.30 | 0.37 | 0.33 | 0.74 |
| **7** | 0.43 | 0.43 | 0.43 | 0.91 |
| **8** | 0.50 | 0.33 | 0.40 | 0.96 |

#### Lesion Score Thresholds for Decisions on Kidney Allocation

*Supplementary Table 8: Final* Remuzzi score used in kidney utilization decisions: Derived from the sum of the four individual lesion scores.

| Score Range | Clinical Implication |
|---|---|
| 0-4 (Mild) | Allocation for transplant |
| 5 (Moderate) | Allocate/Dual Kidney transplantation/Discard (discretion of physician) |
| 6-12 (Severe) | Discard/Consider dual kidney transplantation |

#### Cohen’s Kappa Reference Values

*Supplementary Table 9:* Cohen’s Kappa statistic reference values for the quantification of the level of agreement between two raters (either pathologists or ComPRePS).

| kappa Value | Reliability |
|---|---|
| 0.00–0.50 | Poor |
| 0.51–0.75 | Moderate |
| 0.76–0.90 | Good |
| >0.90 | Excellent |

#### Clinical Use-case 2 Results: Quantitative Morphological Analysis for Kidney Diseases: A study in AN, DN and MCD cohort

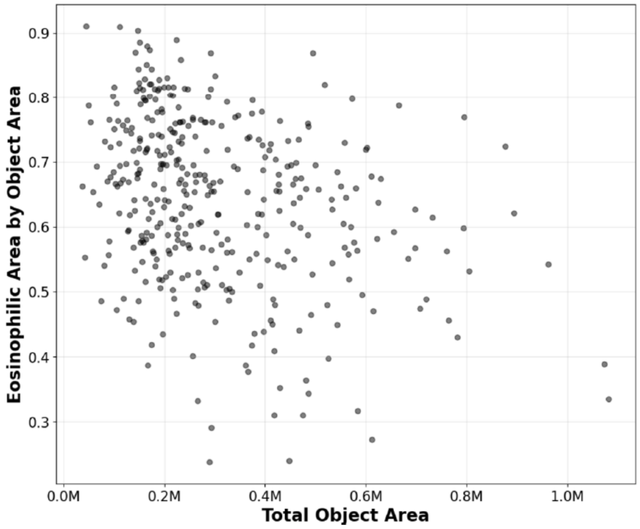


(a) Eosinophilic area by object area vs. total object area

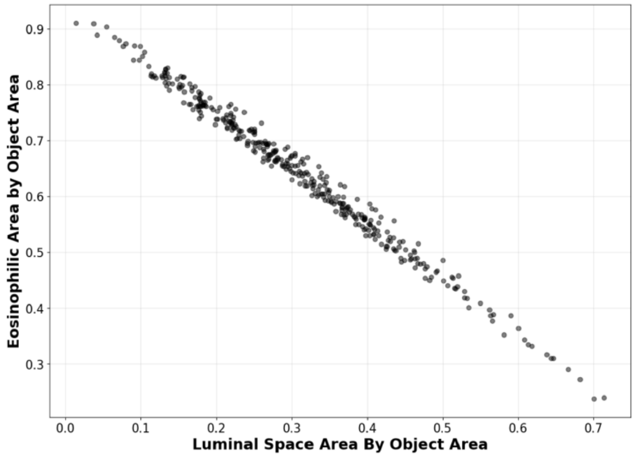


*Supplementary Figure 1:* Scatter plots comparing glomerular features

### Supplementary Methods

#### Plugins Available in ComPRePS

##### *Multi-Compartment Segmentation*

We recently developed [1] a panoptic segmentation model to detect and segment viable and sclerotic glomeruli, cortical and medullary interstitia, tubules, and arteries/arterioles in digitized images from periodic acid–Schiff-stained human nephrectomy sections. Here, we integrate popular Detectron2 [1] panoptic segmentation network within ComPRePS and deploy it as part of our suite of plugins. The segmentation algorithm is packaged as a Docker container using Docker software (Docker Inc., Palo Alto, CA), following Slicer CLI workflow. This multi-compartment segmentation plugin effectively runs the Detectron2 network, segments six micro-anatomical structures and posts the annotations in JSON format for display as annotation layers in the UI as shown in Figure 6 (zoomed views of segmented Functional Tissue Unit (FTU)s can be seen in Figures 7-12). The multi-compartment segmentation codebase is publicly available via GitHub [1] and is easily accessible to the digital pathology community.

##### *Interstitial Fibrosis Tubular Atrophy (IFTA) Segmentation*

Interstitial fibrosis and tubular atrophy (IFTA) are primary indicators of irrecoverable kidney injury. IFTA assessment is conventionally performed visually by a pathologist, where overall percentage of scarred versus unscarred renal cortex is estimated. This process can vary between pathologists due to differences in each pathologist's assessment of IFTA [2]. There's a growing body of literature on the use of digital image analysis, and robust machine learning methods have also been developed to accurately identify and quantify IFTA in WSIs [3]. Recently, we introduced a robust CNN-based IFTA detection model, which demonstrated a statistically significant correlation with all patient-outcome variables [2]. Here, we include the IFTA segmentation pipeline as part of the suite of plugins that we developed here for wider community use. The IFTA plugin is available under ComPRePS. We also provide a trained IFTA model, trained on a set of 79 slides coming from five different institutes. The training set consists of 63 native kidney biopsy specimens, including cases of diabetic nephropathy, 38 transplant kidney biopsy specimens, 13 deceased donor kidney biopsy specimens, and three nephrectomy samples from parenchyma uninvolved by renal cell carcinoma.

##### *Peritubular Capillaries (PTC) Segmentation*

Peritubular capillaries (PTCs) supply oxygen and nutrients to tubular and interstitial cells of the kidney cortex and are thus critical to their integrity. They also provide a route of migration of immune cells to reach tubules and interstitium, giving rise to acute injury. Capillary injury can ultimately lead to an irreversible loss of PTC and is a major indicator of chronic kidney disease (CKD) [4]. One significant feature of ComPRePS is its interoperability, bringing tools and pipelines developed in different laboratories under one unified platform. Here, as part of interoperability, we integrated the pipeline developed by Chen *et al.* [5][29], shared by Dr. Laura Barisoni (Professor, Duke), and implemented PTC segmentation as a plugin as part of the ComPRePS plugin suites (see plugin output in Figure 13). In addition, the trained model used in [6] is also available under M*odel Zoo* in our platform.

##### *Podocyte Detection*

Podocytes are highly specialized epithelial cells that maintain the kidney filtration barrier. It is hypothesized that podocyte loss in the setting of podocytopathic injury from hyperfiltration, hyperglycemia, or hypertension, is an early determinant of proteinuria and glomerulosclerosis across various chronic kidney diseases [7-9]. Therefore, the detection and enumeration of podocytes offer a measurable indicator of irreversible glomerular injury and therapeutic success in these conditions. We have previously developed an automated podometric tool for single section podocyte estimation from WSIs, namely PodoCount [9], with the aim of evaluating WSIs of kidney sections and biopsies immunostained with a podocyte marker through stain deconvolution and local mean-based thresholding techniques. In addition, this tool was also used to study end-stage kidney disease (ESKD) outcomes of a human Diabetic Nephropathy (DN) cohort. The segmentation performance of the developed model was evaluated by computing sensitivity, specificity, precision, and accuracy per glomerulus using a Hit-Miss approach. High performance was observed with near perfect scores for each metric. As part of ComPRePS plugin suite, we develop PodoCount to enable users to detect and quantify podocyte nuclei directly through the web interface.

##### *Arterial Intimal Fibrosis Computation*

Under the ComPRePS plugin suites, we offer an arterial intimal fibrosis computation pipeline [10]. The plugin calculates the intimal area and intimal thickness. The segmentation model is built as an attention U-NET which uses Attention Gates combined with the Residual skip connections.

The inference plugin detects all possible intima annotations, processes and crops the arteries with a fixed margin, uses the model to generate predictions for these cropped images, and performs calculations on the predictions to estimate intimal thickness and stenosis ratio. The plugin saves all cropped artery images, predictions. and logs the required calculation for each image. In HistomicUI web interface, users can browse through the artery crops, visualize the images and predictions, and observe the intimal thickening.

##### *Image Registration*

Medical images are scanned and acquired using diverse imaging technologies, producing data in a variety of formats and modalities. For precise cross-modality comparisons and integrative analysis, the images must be spatially aligned. The process of transforming images into a common coordinate system is called image registration. To support this process, ComPRePS offers user-friendly registration plugins that enable practical and efficient image alignment.

ComPRePS plugins include an image registration plugin [11] that can align two images uploaded to the platform and offers two options rigid body registration which performs intensity-based registration and affine transform which leverages landmark point annotations that are provided for both images. The output registered images are automatically uploaded in the designated folder for review and can be used for downstream analysis. In addition to these linear methods, ComPRePS also offers a thin plate spline (TPS) transformation-based registration method that is able to capture non-linear deformations in images. The method relies on a set of landmarks (or reference) points annotated on an image pair. Unlike rigid or affine transformations, which apply a global transformation across the entire image, TPS can capture local non-linear deformations more effectively. This makes it particularly useful for handling artifacts such as tissue tears, missing regions, and local distortions commonly encountered in rapidly evolving spatial imaging technologies.

##### *Interoperability: Glomeruli Segmentation Models*

Our ComPRePS platform allows running pipelines developed by other labs. With minimal reorganization of code developed by other researchers, these pipelines can be run as containerized software following the Slicer CLI workflow. Currently, there are two glomeruli segmentation models integrated into our platform that were developed by other scientists.

The first plugin is for glomeruli segmentation in frozen kidney tissue sections [12] developed by Duke and Emory University. A nine-layer convolutional neural network, based on the common U-Net architecture, was developed and trained on 258 whole slide images (WSI) from cadaver donor kidney biopsies. ComPRePS currently hosts the prediction pipeline based on this trained model.

The second plugin, developed by Jain *et al.* [13], originates from the Hacking the Human Body competition hosted by the Human Biomolecular Atlas (HuBMAP) and the Human Protein Atlas (HPA) on the Kaggle platform. It leverages the SegFormer mit-b4 model and trained on a dataset of 880 images, containing data from HPA and HuBMAP. The model achieved a Dice score of 0.96 for glomeruli segmentation in periodic acid-Schiff (PAS) stained images of kidney tissues. The inference model is available as a plugin under ComPRePS.

#### Combined Feature Extraction

Quantification of morphometrics from segmented structures is crucial for understanding and quantifying disease states, progression, and prognosis [14]. Within ComPRePS, we developed a variety of feature extraction tools to automate the measurement of kidney histomorphometric data at different levels of granularity. Existing morphometrics were often measured through labor-intensive methods, and automation is essential for obtaining derived data in an objective and robust fashion.

After segmentation, the feature extraction pipeline derives pathologist-informed morphometric features from segmented FTUs at varying levels of granularity. ComPRePS organizes these Pathomic features from coarse to fine detail into three categories: Explainable, Extended clinical and Expanded granular [14, 15]. At the high-level, the pipeline uses image analysis techniques to extract image and contour-based pathomic features including, area, mesangial area for each glomerulus, average tubular basement membrane (TBM) thickness, average cell thickness, luminal fraction for each tubule, and arterial area for each artery. The extended clinical features also include area and radius of each FTU. The expanded granular plugin performs sub-compartmentalization of kidney microanatomical structures into nuclei, Eosinophilic regions and Lumens. It then extracts features including morphology, texture, color and distance transformation.

#### Feature Visualization

In ComPRePS UI, the users can visualize measured features from segmented tissues by the following steps: First, by selecting a whole slide or by circle a region in the slide and see the aggregate statistics from the extracted features highlighted in the lower panel. Second, select up to three regions and visualize difference in features across slides (up to a maximum of ten slides) (shown in Supplementary Figure 1); and finally in the violin plots, when the user hovers over a dot, it highlights the corresponding region as illustrated in the lower panel of Supplementary Figure 2.

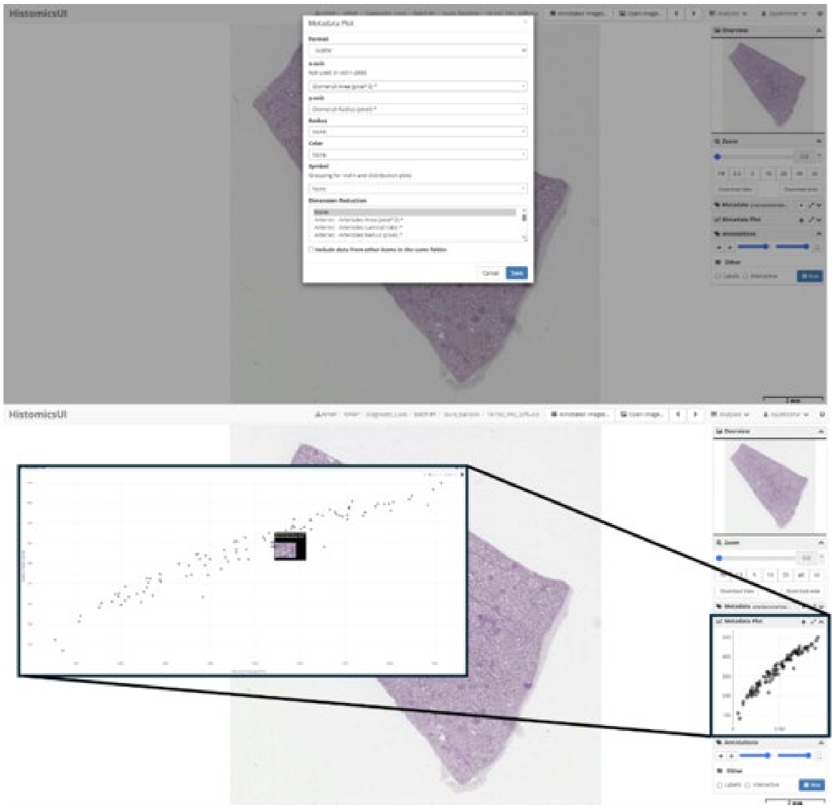

*Supplementary Figure 2:* Feature Visualization in the UI. The users can select the measured features from the segmented FTU. The desired feature columns can be selected in the metadata plot section of the UI as shown in the upper panel. In addition to feature data for the selected image, the users can include data from other images in the same folder for cross slide comparison. The scatter plot of the data selected image is shown in the lower panel. The users can also visualize a particular FTU by hovering their mouse on the points.

#### Available Models for Use in Our Model Zoo

*Supplementary Table 10:* Current Models in the *Model Zoo*.

| **Plugin** | **Developers** | **Purpose** |
|---|---|---|
| Multi-Compartment Segmentation | Lucarelli, N. *et al.*, Kidney360, 2023 | Segment kidney WSI into 6 compartments, namely: cortical interstitium, medullary interstitium, non-sclerotic glomerulus, sclerotic glomerulus, tubule, and artery/arteriole. |
| Peritubular capillaries (PTC) Segmentation | Chen, Y. *et al.*, Kidney360, 2023 | Segment the Peritubular Capillaries in Kidney |
| IFTA and glomerulosclerosis segmentation | Ginley, B. *et al.*, JASN, 2021 | Computational Detection of Interstitial Fibrosis, Tubular Atrophy, and Glomerulosclerosis in Kidney |
| Frozen Glomeruli Segmentation | Li, X. *et al.* JMI, 2021 | Segmentation of glomeruli on kidney donor frozen sections |
| PAS Glomeruli Segmentation | Jain, Y. *et al.*, Nat Communications, 2023 | Segmentation of glomeruli on kidney donors based on PAS staining |
| PodoCount | Briana, S. *et al.*, Kidney International Reports, 2022 | A Robust, Fully Automated, Whole-Slide Podocyte Quantification Tool |
| DeepCell | Moen, E. *et al.*, Nat Methods, 2019 | Deep Learning library for single-cell analysis of biological images |

#### Data Services

WSIs are typically very large, ranging from hundreds of megabytes to tens of gigabytes. Consequently, efficient data storage and management are critical components of any computational pathology platform. Moreover, since researchers and pathologists frequently work with Protected Health Information (PHI) [16], it is imperative that the storage infrastructure adheres to stringent security standards to protect this sensitive data.

ComPRePS addresses these challenges through its integration with HiPerGator. Specifically, ComPRePS leverages HiperGator's Lustre-based [17, 18] parallel file system— a network-based system ideally suited for managing large-scale data. This file system enables users to access and share data without unnecessary duplication, thus optimizing both storage space and data retrieval speeds.

HiperGator currently offers three distinct tiers of storage - Red, Blue, and Orange - to support varying performance and storage needs. The Red tier provides high-performance storage, designed for projects that demand the highest input/output (I/O) performance. However, for most applications, the Blue and Orange tiers are more commonly used. The Blue tier offers a high-performance parallel file system optimized for frequent I/O operations, while the Orange tier, although slower, is well-suited for long-term storage needs. Users can upload their data into any of these partitions and import it into ComPRePS via a symbolic link. Users must grant read and write access to the linked folder, allowing ComPRePS to retrieve the data for analysis and subsequently export the results back to the original storage location for further downstream analysis. This flexible way data handling ensures that users are not confined to our platform for pre- or post-processing tasks, enabling them to bring their own data into the platform or extract the AI generated results as needed.

Collaboration across multiple institutions is further facilitated by the ComPRePS architecture. A common tool for secure inter-institutional data sharing is Globus [19], which supports fast and secure file transfers. Typically, shared data resides on network-based file systems such as Blue or Orange tiers of the platform. Once the data is prepared, a Globus link can be generated to grant read and/or write access to external collaborators. Since ComPRePS can symbolically link to these file systems, users can begin their analyses immediately following data transfer and can share results as soon as they become available, thereby supporting efficient and seamless collaboration across institutions.

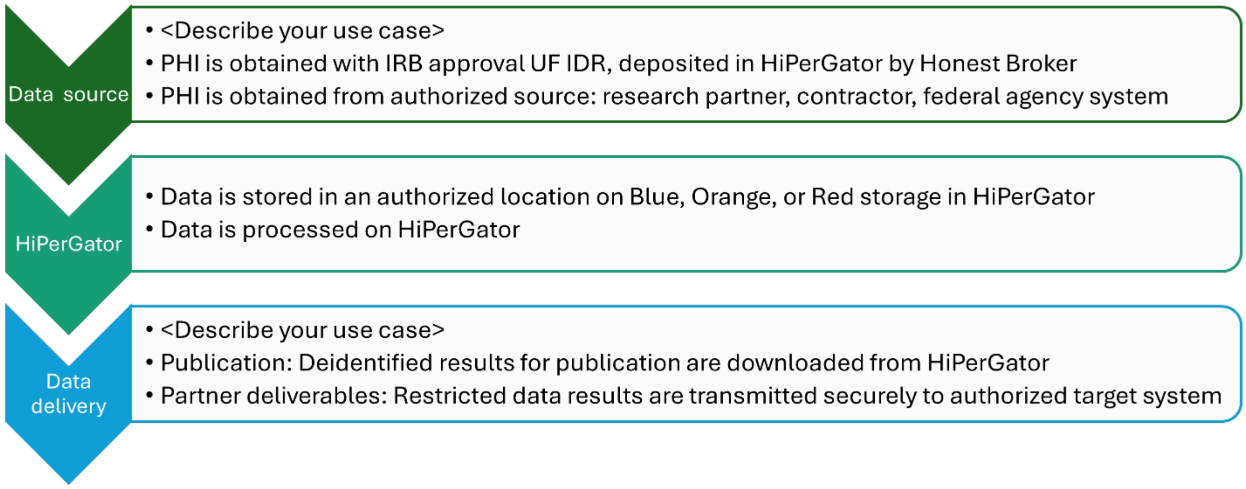


*Supplementary Figure 3:* Data flow diagram depicting UF's protocol for PHI data. *Source: rc.ufl.edu*

For PHI data, HiPerGator provides additional flexibility by allowing storage on any of the three tiers—Blue, Orange, or Red— while ensuring compliance with data security protocols. However, due to the sensitive nature of PHI data, a formal onboarding process must be completed. A flowchart outlining this procedure is shown in Supplementary Figure 2, and further details can be found in [20, 21].

In summary, by integrating advanced storage solutions and secure data transfer protocols, ComPRePS not only addresses the technical challenges associated with large-scale WSI management but also ensures that sensitive PHI data is handled in full compliance with regulatory standards. This robust data management framework is essential for supporting both the computational demands of modern pathology research and the collaborative workflows that drive clinical innovation.

#### Security Implementation

ComPRePS is designed as a secure platform, ensuring robust protection of sensitive PHI (Protected Health Information) data. This section details the security architecture of ComPRePS, outlining the multiple layers of authentication and access control mechanisms that are in place.

First, the platform provides a standard user interface for login and registration. To sign up, users must submit a registration form, which is reviewed and approved by an internal team member. While this is a standard security practice, it is not the primary element of ComPRePS's security model.

The primary security of ComPRePS is enforced through the HiperGator Infrastructure, managed by the University of Florida's Research Computing (UFRC) department. This ensures that the application adheres to the highest security standards.

Access to ComPRePS requires authentication through one of two methods: GatorLink Authentication: Users affiliated with UF already have a GatorLink account, which is part of the university's Identity and Access Management (IAM) system. For external users, a GatorLink account can be created by submitting a request form with the necessary contact details. Once verified by UFIT, an account is provisioned. After obtaining a GatorLink account, users must then apply for HiperGator access, specifying a Principal Investigator (PI) who will sponsor their account. This step prevents unauthorized access and ensures responsible usage of computing resources.

InCommon Federated Login System: This option provides secure single sign-on (SSO) and is widely adopted by educational and research institutions. Users whose organizations participate in the InCommon Federation and have an active institutional affiliation can bypass the GatorLink account creation step and directly request a Federated HiperGator account. Upon approval, they can access the application by initiating a session through the UFApps OnDemand portal, requesting a HiperGator Desktop session, and launching the ComPRePS application from within the browser.

In addition to authentication mechanisms, ComPRePS employs Virtual Private Network (VPN) restrictions, ensuring that the platform can only be accessed from within the UF network. Users must be connected to the UF network via VPN, with different VPN solutions available based on account type: GatorLink users utilize the Cisco Secure Client VPN, whereas non-GatorLink users can connect using eduVPN. These security measures collectively establish ComPRePS as a highly secure platform- particularly crucial when handling PHI and other sensitive research data.

#### Educational and Clinical Utility of ComPRePS

ComPRePS has demonstrated value as an educational and clinical tool in nephropathology. User LR, a clinical nephrologist and frequent user, reported that the platform enhanced educational sessions with pathology trainees by providing intuitive visualizations and quantitative support via segmentation masks. LR noted that the AI-driven histomorphometric features made complex pathological concepts more accessible and highlighted the usefulness of structured slide exploration for rapid identification of renal structures. He also appreciated the platform's responsive slide-viewing experience (when internet connection was stable) but stressed that careful review of AI-generated segmentation was still necessary. LR suggested usability improvements, including simplified toolbars, better control visibility, and an “undo” function for annotation and QC tasks. Overall, LR saw ComPRePS as an effective tool for remote collaboration and improving reproducibility in multicenter studies.

User BF, an independent clinical nephropathologist from University of Coimbra, used ComPRePS as a complementary tool in clinical nephropathology. User BF emphasized the platform’s ability to consistently identify structures that may be missed due to human error or observer fatigue and noted its potential to reduce interobserver variability. She also highlighted ComPRePS’s capacity to extract quantitative features (such as glomerular dimensions and vascular changes) that are difficult to measure manually, which could support new histological endpoints in research. User BF pointed out the importance of minimizing annotator bias and recommended large, well-annotated datasets. She also raised concerns about generalizability due to differences in staining and scanning and recommended integrating clinical variables to enhance interpretability.

User CP, a senior nephrologist and faculty at University of Coimbra, confirmed ComPRePS’s utility for standardizing detection and quantification of renal histological features and reducing interobserver variability. User CP praised the platform’s ability to quickly and accurately segment renal structures but found the manual editing tool cumbersome. Suggested improvements included automatic calculation of key metrics (e.g., percentage of sclerotic glomeruli, interstitial fibrosis, inflammatory cell density) and automatic flagging of abnormal regions for prioritized review.

In summary, real-world feedback from these nephropathologists highlight ComPRePS’s value in clinical education and research. The suggestions provided offer clear guidance for ongoing platform development.

#### Bringing Your AI Model and Data to the ComPRePS Platform

In this section, we outline how to integrate AI models and data into the ComPRePS platform and utilize its capabilities for computational workflows, research collaboration, and clinical insight generation. The typical workflow is depicted in Figure 1 in the main text.

If you have developed an AI model for tasks such as segmentation, to deploy this model on the ComPRePS platform, it must first be converted into a plugin. This plugin should adhere to the publicly available Slicer CLI execution [22] format. Our team provides support for converting AI models into a format compatible with ComPRePS. The slicer_cli_model format packages the code into a containerized image. Currently, the platform utilizes Docker for containerization to maintain backward compatibility with existing plugins and for ease of distribution. However, the future roadmap includes support for directly adding plugins as .sif (Apptainer) images.

An excellent demonstration of the platform’s capabilities is its integration with MONAI [23] (Medical Open Network for AI) pipelines. MONAI provides a collection of cutting-edge deep learning models for medical imaging tasks such as segmentation, annotation, and classification. ComPRePS addresses key challenges commonly encountered when using MONAI, particularly the absence of a user interface (UI) and the requirement for extensive coding environment setup. Through integration with ComPRePS, MONAI pipelines can be executed seamlessly, eliminating the need for complex setup and enabling faster deployment and broader accessibility.

To ensure such integrations work efficiently, each plugin, including MONAI pipelines or other AI models, must expose its capabilities using the –list-cli command. This helps the platform identify and run the relevant functionality during runtime. Once the plugin is added, it can be executed on either user-uploaded data or datasets available publicly on the platform.

An important feature of ComPRePS is its ability to facilitate collaboration. For instance, collaborators from other institutions can grant controlled access to their data, enabling you to run pipelines and collect results. The platform offers flexible data access controls, allowing users to choose appropriate access levels. Conversely, you may allow others to use your model, receiving valuable feedback to refine and enhance its capabilities.

Finally, ComPRePS allows users to export results for independent analysis outside the platform. This enables downstream use of AI outputs in external tools or analyses beyond the ComPRePS ecosystem.

#### Flexibility and Portability of ComPRePS Application Architecture

As detailed in the Methods section, the ComPRePS application architecture is composed of modular, independently functioning components. This modularity, combined with its open-source foundation, gives ComPRePS robust adaptability across various computational environments. Figure 7 of the main text illustrates the core components that constitute the primary functionality of the application. The front-end services can leverage the resources where the application is setup. This would typically be in a virtual machine (VM). Meanwhile, the backend services constitute the computational core of the application and these can leverage any HPC based infrastructure with a SLURM (Simple Linux Utility for Resource Management) interface.

The modularity of ComPRePS allows for independent modification in each component to meet the specific requirements of diverse user environments. For instance, users can easily switch to network-based storage solutions, depending on specific needs and resource availability. Likewise, users without access to high-performance computing (HPC) can deploy ComPRePS on local machines using a monolithic architecture. This setup enables the application to run seamlessly on a single node or edge device without modification, as the application was originally designed with monolithic architectures in mind.

In contrast, for users with access to advanced computing infrastructure, such as the HiperGator (HPG) Supercomputer, ComPRePS can be scaled to accelerate AI computations using parallel and distributed computing frameworks. Deployment on HiperGator required adapting the Worker and Slice-CLI-Web components to support distributed computing. These components were configured to communicate with HPG via SLURM, enabling efficient, on-demand orchestration of tasks across multiple nodes.

This flexibility allows users to configure and install ComPRePS to suit their operational environment - whether a standalone monolithic setup or a distributed HPC system. Additionally, users facing data privacy restrictions (e.g., IRB constraints) can fully deploy ComPRePS within private infrastructure, ensuring compliance with privacy regulations.

#### Feature Visualization

##### *Visualization of Extracted Features in the Metadata Plot*

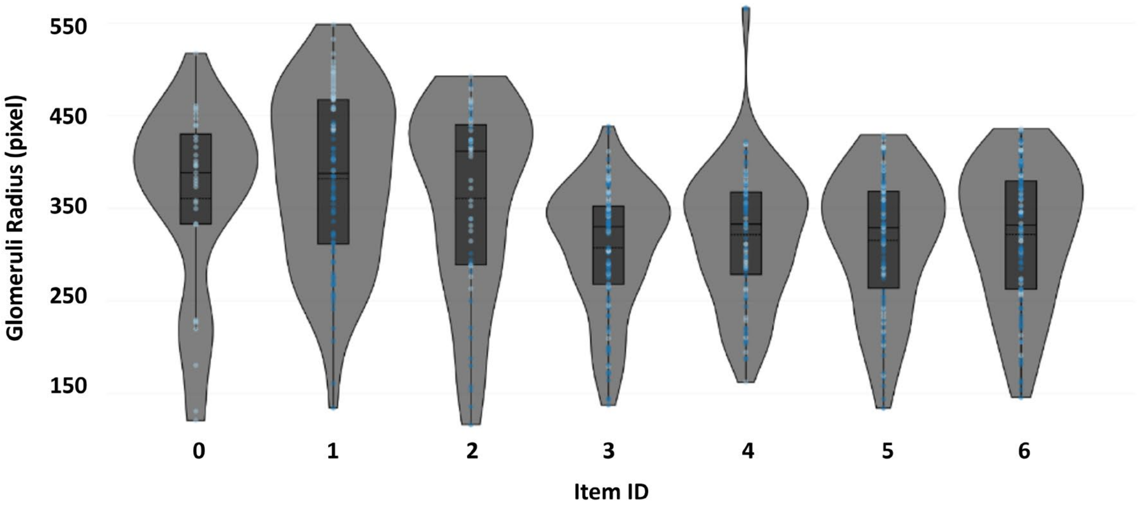


*Supplementary Figure 4:* Inter and intra slide feature visualization. Data distribution is shown as violin plots derived from different slides shown on the upper panel. The tool allows users to select regions in the scatter plot and generate violin plots for the regions.

##### *Feature Visualization Plugin*

The feature visualization plugin enables users to perform rapid visualization and statistical analysis on extracted features from FTUs. Currently, the plugin supports analysis of glomeruli, tubules, and arteries/arterioles, but it is designed to be extensible for additional use cases.

Users can leverage the plugin for rapid visual exploration of extracted features. The plugin provides an option to select a dimensionality reduction algorithm, with two widely used methods available: Principal Component Analysis (PCA) and Uniform Manifold Approximation and Projection (UMAP) [24, 25]. By default, PCA is selected. Users can also select a clustering algorithm—either K-means or Louvain—with K-means set as the default.

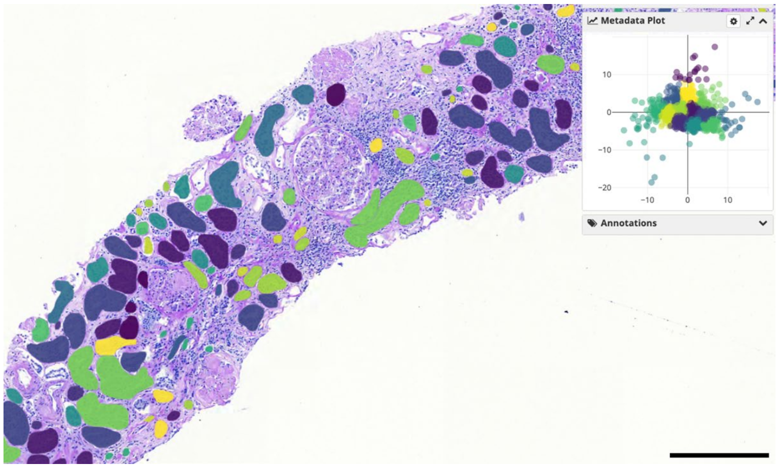


*Supplementary Figure 5:* Feature Visualization of extracted features from Tubules. The scalebar corresponds to 300 μm.

Upon execution, the plugin generates two output files. The first is an annotation layer file, which enables users to visualize the spatial distribution of clusters on the Whole Slide Image (WSI). The second output integrates with the Metadata Plot tool, enabling visualization of cluster distribution within the principal component space. Clusters in the metadata plot share the same colormap as those in the annotation layer, enhancing interpretability of feature distribution (Supplementary Figure 5).

Additionally, users may opt to perform statistical analysis during plugin execution. If selected, the plugin executes the Wilcoxon rank-sum test on the extracted features, generating an Excel file containing the statistical results for further analysis.

##### *Metadata*

Metadata serves as supplementary information that can be associated with histological data in ComPRePS, offering one of the platform's most powerful capabilities. By leveraging metadata, users can perform advanced visualizations, track cohort-specific details at the level of individual slides and enable further downstream analyses and collaborations.

Metadata can be added at multiple levels of granularity by both users and AI algorithms. At the most granular level, metadata can be assigned to each individual Whole Slide Image (WSI). When reviewing a slide in ComPRePS, users can access the Metadata subfield. Clicking on the plus icon allows users to manually add metadata in various formats, including JSON, text, or predefined fields such as Glomerular QC, Participant ID, and other cohort-specific attributes. Additionally, metadata at the slide level can be automatically generated by AI algorithms, facilitating both individual WSI analysis and cohort-level comparisons.

For handling larger datasets, metadata can also be added programmatically. This allows users to upload extensive metadata efficiently by writing a simple script and leveraging the CompRePS API, rather than entering data manually. Alternatively, users may upload metadata via an Excel file directly through the UI.

To ensure compatibility with the Metadata Plot visualization tool, users must format the Excel metadata files accordingly. Specifically, files must include bounding box (bbox) values—xmin, xmax, ymin, ymax—for the relevant features. This formatting is only required if users intend to utilize the plotting tool; otherwise, it is optional.

ComPRePS offers extensive flexibility for managing metadata, whether created within the platform or imported from external systems. Regardless of the method used for metadata integration, users can seamlessly access and utilize this information to enhance their analyses.

*Supplementary Table 11:* List of features extracted from Glomeruli. Abbreviations: D - Distance Transform features, C - Color features, T - Texture features and M - Morphological features

| Index | Type | Name |
|---|---|---|
| 1 | D | Max Distance Transform By Object Area Luminal Space |
| 2 | D | Max Distance Transform By Object Area Eosinophilic |
| 3 | D | Max Distance Transform By Eosinophilic Area |
| 4 | D | Max Distance Transform Eosinophilic |
| 5 | D | Sum Distance Transform By Nuclei Area |
| 6 | D | Mean Distance Transform By Eosinophilic Area |
| 7 | D | Sum Distance Transform By Object Area Eosinophilic |
| 8 | D | Max Distance Transform Nuclei |
| 9 | D | Sum Distance Transform Luminal Space |
| 10 | D | Mean Distance Transform Luminal Space |
| 11 | D | Sum Distance Transform By Luminal Space Area |
| 12 | D | Max Distance Transform By Nuclei Area |
| 13 | D | Mean Distance Transform Eosinophilic |
| 14 | D | Mean Distance Transform By Object Area Eosinophilic |
| 15 | D | Sum Distance Transform By Eosinophilic Area |
| 16 | D | Sum Distance Transform Nuclei |
| 17 | D | Sum Distance Transform By Object Area Luminal Space |
| 18 | D | Max Distance Transform By Object Area Nuclei |
| 19 | D | Sum Distance Transform Eosinophilic |
| 20 | D | Mean Distance Transform By Luminal Space Area |
| 21 | D | Mean Distance Transform By Object Area Nuclei |
| 22 | D | Mean Distance Transform Nuclei |
| 23 | D | Mean Distance Transform By Object Area Luminal Space |
| 24 | D | Max Distance Transform By Luminal Space Area |
| 25 | D | Max Distance Transform Luminal Space |
| 26 | D | Mean Distance Transform By Nuclei Area |
| 27 | D | Sum Distance Transform By Object Area Nuclei |
| 28 | C | Mean Green Luminal Space |
| 29 | C | Standard Deviation Blue Luminal Space |
| 30 | C | Standard Deviation Red Nuclei |

| | | |
|---|---|---|
| 31 | C | Mean Blue Nuclei |
| 32 | C | Standard Deviation Green Luminal Space |
| 33 | C | Mean Red Eosinophilic |
| 34 | C | Mean Green Eosinophilic |
| 35 | C | Mean Red Nuclei |
| 36 | C | Mean Green Nuclei |
| 37 | C | Mean Blue Eosinophilic |
| 38 | C | Standard Deviation Green Eosinophilic |
| 39 | C | Standard Deviation Blue Eosinophilic |
| 40 | C | Standard Deviation Red Eosinophilic |
| 41 | C | Standard Deviation Red Luminal Space |
| 42 | C | Standard Deviation Green Nuclei |
| 43 | C | Standard Deviation Blue Nuclei |
| 44 | C | Mean Red Luminal Space |
| 45 | C | Mean Blue Luminal Space |
| 46 | T | Homogeneity Nuclei |
| 47 | T | Energy Eosinophilic |
| 48 | T | Contrast Nuclei |
| 49 | T | Correlation Nuclei |
| 50 | T | Correlation Eosinophilic |
| 51 | T | Homogeneity Luminal Space |
| 52 | T | Correlation Luminal Space |
| 53 | T | Contrast Eosinophilic |
| 54 | T | Contrast Luminal Space |
| 55 | T | Energy Luminal Space |
| 56 | T | Homogeneity Eosinophilic |
| 57 | T | Energy Nuclei |
| 58 | M | Eosinophilic Area |
| 59 | M | Major Axis Length |
| 60 | M | Nuclei Area By Object Area |
| 61 | M | Luminal Space Area |
| 62 | M | Luminal Space Area By Object Area |
| 63 | M | Nuclei Number |
| 64 | M | Total Object Perimeter |
| 65 | M | Total Object Aspect Ratio |
| 66 | M | Nuclei Area |

| | | |
|---|---|---|
| 67 | M | Mean Nuclear Area |
| 68 | M | Eosinophilic Area By Object Area |
| 69 | M | Standard Deviation Aspect Ratio Nuclei |
| 70 | M | Total Object Area |
| 71 | M | Mean Aspect Ratio Nuclei |
| 72 | M | Minor Axis Length |
| 73 | M | Mesangial Area |
| 74 | M | Mesangial Fraction |
| 75 | M | Area |
| 76 | M | Radius |

*Supplementary Table 12:* List of features extracted from Arteries/Arterioles. Abbreviations: D - Distance Transform features, C - Color features, T - Texture features, M - Morphological features

| Index | Type | Name |
|---|---|---|
| 1 | D | Mean Distance Transform By Object Area Nuclei |
| 2 | D | Max Distance Transform Eosinophilic |
| 3 | D | Max Distance Transform By Object Area Nuclei |
| 4 | D | Sum Distance Transform By Nuclei Area |
| 5 | D | Mean Distance Transform Luminal Space |
| 6 | D | Sum Distance Transform By Eosinophilic Area |
| 7 | D | Max Distance Transform Luminal Space |
| 8 | D | Mean Distance Transform By Object Area Eosinophilic |
| 9 | D | Max Distance Transform By Eosinophilic Area |
| 10 | D | Mean Distance Transform Eosinophilic |
| 11 | D | Max Distance Transform By Object Area Eosinophilic |
| 12 | D | Sum Distance Transform Nuclei |
| 13 | D | Max Distance Transform Nuclei |
| 14 | D | Max Distance Transform By Nuclei Area |
| 15 | D | Mean Distance Transform By Luminal Space Area |
| 16 | D | Sum Distance Transform Luminal Space |
| 17 | D | Mean Distance Transform By Object Area Luminal Space |
| 18 | D | Sum Distance Transform Eosinophilic |
| 19 | D | Max Distance Transform By Object Area Luminal Space |
| 20 | D | Sum Distance Transform By Object Area Luminal Space |
| 21 | D | Sum Distance Transform By Object Area Nuclei |
| 22 | D | Sum Distance Transform By Object Area Eosinophilic |

| | | |
|---|---|---|
| 23 | D | Max Distance Transform By Luminal Space Area |
| 24 | D | Sum Distance Transform By Luminal Space Area |
| 25 | D | Mean Distance Transform By Nuclei Area |
| 26 | D | Mean Distance Transform Nuclei |
| 27 | D | Mean Distance Transform By Eosinophilic Area |
| 28 | C | Mean Blue Nuclei |
| 29 | C | Standard Deviation Blue Luminal Space |
| 30 | C | Standard Deviation Red Eosinophilic |
| 31 | C | Standard Deviation Blue Eosinophilic |
| 32 | C | Standard Deviation Red Nuclei |
| 33 | C | Standard Deviation Green Eosinophilic |
| 34 | C | Mean Green Eosinophilic |
| 35 | C | Mean Red Luminal Space |
| 36 | C | Standard Deviation Green Nuclei |
| 37 | C | Standard Deviation Green Luminal Space |
| 38 | C | Mean Blue Eosinophilic |
| 39 | C | Mean Red Eosinophilic |
| 40 | C | Mean Blue Luminal Space |
| 41 | C | Mean Green Nuclei |
| 42 | C | Mean Green Luminal Space |
| 43 | C | Standard Deviation Red Luminal Space |
| 44 | C | Standard Deviation Blue Nuclei |
| 45 | C | Mean Red Nuclei |
| 46 | T | Energy Luminal Space |
| 47 | T | Contrast Eosinophilic |
| 48 | T | Homogeneity Nuclei |
| 49 | T | Correlation Luminal Space |
| 50 | T | Homogeneity Luminal Space |
| 51 | T | Energy Nuclei |
| 52 | T | Contrast Luminal Space |
| 53 | T | Contrast Nuclei |
| 54 | T | Energy Eosinophilic |
| 55 | T | Correlation Eosinophilic |
| 56 | T | Homogeneity Eosinophilic |
| 57 | T | Correlation Nuclei |
| 58 | M | Luminal Space Area By Object Area |

| | | |
|---|---|---|
| 59 | M | Total Object Area |
| 60 | M | Mean Nuclear Area |
| 61 | M | Eosinophilic Area By Object Area |
| 62 | M | Nuclei Number |
| 63 | M | Total Object Perimeter |
| 64 | M | Total Object Aspect Ratio |
| 65 | M | Major Axis Length |
| 66 | M | Standard Deviation Aspect Ratio Nuclei |
| 67 | M | Mean Aspect Ratio Nuclei |
| 68 | M | Luminal Space Area |
| 69 | M | Eosinophilic Area |
| 70 | M | Nuclei Area By Object Area |
| 71 | M | Minor Axis Length |
| 72 | M | Nuclei Area |
| 73 | M | Arterial Area |
| 74 | M | Radius |
| 75 | M | Luminal Ratio |

*Supplementary Table 13.* List of features extracted from Tubules. Abbreviations: D - Distance Transform features, C - Color features, T - Texture features, M - Morphological features, S - Spatial features

| Index | Type | Name |
|---|---|---|
| 1 | D | Sum Distance Transform By Luminal Space Area |
| 2 | D | Mean Distance Transform By Luminal Space Area |
| 3 | D | Sum Distance Transform By Object Area Eosinophilic |
| 4 | D | Max Distance Transform By Object Area Luminal Space |
| 5 | D | Sum Distance Transform By Object Area Nuclei |
| 6 | D | Max Distance Transform By Object Area Eosinophilic |
| 7 | D | Sum Distance Transform By Object Area Luminal Space |
| 8 | D | Max Distance Transform Luminal Space |
| 9 | D | Max Distance Transform Eosinophilic |
| 10 | D | Max Distance Transform By Nuclei Area |
| 11 | D | Mean Distance Transform By Object Area Eosinophilic |
| 12 | D | Sum Distance Transform Luminal Space |
| 13 | D | Mean Distance Transform By Object Area Nuclei |
| 14 | D | Mean Distance Transform By Eosinophilic Area |

| | | |
|---|---|---|
| 15 | D | Max Distance Transform By Eosinophilic Area |
| 16 | D | Mean Distance Transform By Object Area Luminal Space |
| 17 | D | Mean Distance Transform Eosinophilic |
| 18 | D | Mean Distance Transform By Nuclei Area |
| 19 | D | Max Distance Transform By Object Area Nuclei |
| 20 | D | Sum Distance Transform Nuclei |
| 21 | D | Sum Distance Transform Eosinophilic |
| 22 | D | Sum Distance Transform By Nuclei Area |
| 23 | D | Max Distance Transform Nuclei |
| 24 | D | Mean Distance Transform Nuclei |
| 25 | D | Max Distance Transform By Luminal Space Area |
| 26 | D | Sum Distance Transform By Eosinophilic Area |
| 27 | D | Mean Distance Transform Luminal Space |
| 28 | C | Mean Green Nuclei |
| 29 | C | Standard Deviation Red Nuclei |
| 30 | C | Standard Deviation Green Eosinophilic |
| 31 | C | Mean Green Luminal Space |
| 32 | C | Mean Red Luminal Space |
| 33 | C | Standard Deviation Blue Eosinophilic |
| 34 | C | Mean Blue Nuclei |
| 35 | C | Mean Blue Luminal Space |
| 36 | C | Mean Red Eosinophilic |
| 37 | C | Standard Deviation Blue Luminal Space |
| 38 | C | Mean Green Eosinophilic |
| 39 | C | Standard Deviation Green Nuclei |
| 40 | C | Standard Deviation Red Luminal Space |
| 41 | C | Mean Blue Eosinophilic |
| 42 | C | Standard Deviation Red Eosinophilic |
| 43 | C | Mean Red Nuclei |
| 44 | C | Standard Deviation Green Luminal Space |
| 45 | C | Standard Deviation Blue Nuclei |
| 46 | T | Correlation Eosinophilic |
| 47 | T | Contrast Nuclei |
| 48 | T | Correlation Luminal Space |
| 49 | T | Energy Nuclei |
| 50 | T | Homogeneity Eosinophilic |

| | | |
|---|---|---|
| 51 | T | Homogeneity Luminal Space |
| 52 | T | Energy Luminal Space |
| 53 | T | Homogeneity Nuclei |
| 54 | T | Contrast Luminal Space |
| 55 | T | Energy Eosinophilic |
| 56 | T | Contrast Eosinophilic |
| 57 | T | Correlation Nuclei |
| 58 | M | Luminal Space Area By Object Area |
| 59 | M | Eosinophilic Area By Object Area |
| 60 | M | Eosinophilic Area |
| 61 | M | Nuclei Area By Object Area |
| 62 | M | Major Axis Length |
| 63 | M | Nuclei Number |
| 64 | M | Mean Nuclear Area |
| 65 | M | Minor Axis Length |
| 66 | M | Nuclei Area |
| 67 | M | Mean Aspect Ratio Nuclei |
| 68 | M | Total Object Aspect Ratio |
| 69 | M | Standard Deviation Aspect Ratio Nuclei |
| 70 | M | Total Object Area |
| 71 | M | Total Object Perimeter |
| 72 | M | Luminal Space Area |
| 73 | M | Average TBM Thickness |
| 74 | M | Average Cell Thickness |
| 75 | M | Luminal Fraction |
| 76 | M | Radius |
| 77 | S | In Medulla |

#### Clinical Use - Case 1 Methods: Donor Biopsy Assessment for Kidney Transplant

##### Rapid Formalin-Fixed Paraffin-Embedded Tissue Protocol

**Fixation-Dehydration:**

Tissue samples are fixed in 10% neutral buffered formalin for 15 minutes, with gentle agitation and temperature control to improve fixation uniformity and minimize autolysis. Following fixation, samples undergo dehydration in 100% alcohol, lasting 10 minutes, 10 minutes, and 20 minutes, respectively, to ensure complete removal of water prior to clearing.

**Clearing:**
Samples are then immersed in xylene for 5, 5, and 10 minutes, under agitation and controlled temperature, to facilitate complete replacement of alcohol with clearing agent and promote optimal paraffin infiltration.

**Embedding:**
After clearing, tissues are embedded in paraffin wax at 62°C for 10 and 20 minutes. Controlled temperature and agitation are maintained throughout this step to preserve tissue architecture and minimize artifact formation.

**Staining:**
Renal tissue sections (2–3 μm thick) are stained with Hematoxylin and Eosin (H&E) and Periodic Acid–Schiff (PAS) stains.

#### Donor Demographic Data

*Supplementary Table 14:* Donor demographic and clinical variables. KDPI, Kidney Donor Profile Index

| Demographic and Clinical variables | All *n*=152 (100%) | Allocated *n*=110 (72.4%) | Discarded *n*=42 (27.6%) | *p* |
|---|---|---|---|---|
| **Gender (male)** | 94 (61.8%) | 71 (64.5%) | 23 (54.8%) | NS |
| **Race (white)** | 152 (100%) | 110 (100%) | 42 (100%) | NS |
| **Age (years)** | 64.0 ± 12.9 | 62.3± 13.6 | 68.7± 9.2 | 0.008 |
| **Cause of donor death** | | | | 0.006 |
| **Stroke** | 113 (74.3%) | 75 (68.2%) | 38 (90.5%) | NS |
| **Anoxia** | 13 (8.5%) | 8 (7.3%) | 4 (9.5%) | NS |
| **DCD** | 1 (0.7%) | 1 (0.9%) | 0 (0%) | NS |
| **Trauma** | 26 (17%) | 26 (23.6%) | 0 (0%) | NS |
| **Height (meters)** | 166.5 ± 6.7 | 167.00 ± 6.1 | 164.3 ± 8.5 | NS |
| **Weight (kilograms)** | 76.7 ± 12.3 | 76.3 ± 13.4 | 78.1 ± 11.5 | NS |
| **Diabetes history** | 17 (11.2%) | 5 (4.5%) | 12 (28.6%) | <0.001 |
| **Hypertension history** | 104 (68.4%) | 73 (66.4%) | 31 (73.8%) | NS |
| **HCV status** | 0 (0%) | 0 (0%) | 0 (0%) | NS |
| **Donor serum creatinine (mg/dL)** | 0.90 ± 0.4 | 0.8 ± 0.3 | 1.1 ± 0.5 | <0.001 |
| **Donor eGFR CKD-EPI 2021 (ml/min/1.732)** | 87.1 ± 22.0 | 91.22 ± 20.0 | 72.2 ± 22.9 | <0.001 |
| **KDPI** | 74.0 ± 21.8 | 69.5 ± 22.6 | 87.2 ± 12.2 | <0.001 |

#### Histopathological Score Calculations

##### Assessment of the Cortical Area of each WSI

We leveraged the ComPRePS multi-compartment segmentation tool to segment the entire tissue section, medullary region, glomeruli, and vascular structures, and to compute the corresponding area for each compartment. All area measurements are displayed within the interface. Using this information, the cortical area was calculated based on the following formula:

- $C_A = T_A - M_A - NSG_A - SG_A - V_A$
  - where:
    - $C_A$: Cortical area
    - $T_A$: Total area
    - $M_A$: Medullary area
    - $NSG_A$: Non-Sclerotic glomerular area
    - $SG_A$: Sclerotic glomerular area
    - $V_A$: Vascular area

##### *Converting Computational IFTA to Remuzzi IF and TA Scores*

Since pathologists score interstitial fibrosis (IF) and tubular atrophy (TA) as separate components, whereas the ComPRePS computational pipeline generates a single unified score for IFTA, we developed a harmonization strategy to align both approaches. Specifically, we combined the pathologist assigned IF and TA scores into a single IFTA score by selecting the higher of the two values and doubling it, provided that either IF or TA was greater than 0. The resulting composite IFTA score followed the grading scale: absent (0 points), <20% (2 points), 20–50% (4 points), and >50% (6 points). This method enabled consistent comparison between human and computational scoring. Since the computational cortical IFTA area is continuous, we adopted a data-derived threshold using $10^{\text{th}}$ the percentile of the calculated IFTA distribution and assigned a zero score for areas <0.006%.

- Confusion matrices comparing ComPRePS predictions with the ground truth (GT), as well as those comparing GT with on-call pathologist assessments.
- Prediction metrics (precision, recall, F1 score, and accuracy) for ComPRePS in relation to on-call pathologists, using the GT as a reference standard.

##### *Converting Computational Lesion Values for Glomerulosclerosis, IFTA and intimal fibrosis*

The individual computational values for glomerulosclerosis, IFTA, and intimal fibrosis were converted to lesion scores using predefined thresholds (Supplementary Table 15) and combined to compute the computational Remuzzi grade.

*Supplementary Table 15:* Histopathological classification for semiquantitative grading of chronic lesions in donor kidneys

| | 0 | 1 | 2 | 3 |
|---|---|---|---|---|
| Global glomerulosclerosis | None | < 20% | 20-50% | > 50% |
| Tubular atrophy | Absent | < 20% | 20-50% | > 50% |
| Interstitial fibrosis | Absent | < 20% | 20-50% | > 50% |
| Arterial intimal fibrosis | Absent | < 25% | 25 - 50% | > 50% |

##### *Remuzzi Score Calculation by On-Call and Reviewer Pathologists*

On-call and reviewing pathologists assessed the histological lesions in each donor biopsy to assign a Remuzzi score, following the approach outlined below:

- Percentage of global glomerulosclerosis: The percentage of glomerulosclerosis was determined by calculating the ratio of globally sclerotic glomeruli to the total number of glomeruli present in the histologic section. The glomerulosclerosis percentage number was subsequently categorized into predefined thresholds: none (0 points), <20% (1 point), 20–50% (2 points), and >50% (3 points).
- Interstitial fibrosis (IF): Pathologists evaluated the percentage of the renal cortical area replaced by fibrous connective tissue and subsequently categorized it as: absent (0 points), <20% (1 point), 20–50% (2 points), or >50% (3 points).
- Tubular atrophy (TA): The extent of tubular atrophy was estimated as the percentage of cortical tubules affected by atrophic changes and categorized as: absent (0 points), <20% (1 point), 20–50% (2 points), or >50% (3 points).
- Arterial intimal fibrosis: A visual analog scale for assessing arterial intimal fibrosis was used, corresponding to the Banff Lesion Score cv for the most severely affected artery. Intimal fibrosis is assessed by measuring fibro-intimal thickening for the most severely affected artery determined by pathologists with the following classes: none (0 points), <25% (1 point), 26–50% (2 points), or >50% vascular narrowing (3 points).

The four lesion scores are then added to generate the final Remuzzi grade.

#### Navigating WSIs with Annotation Visualizations through ComPRePS UI

Users can access the Computational Microscopy Imaging Lab Slide Archive (https://athena.rc.ufl.edu/) as a public user (username: public, password: public, Supplementary Figure 6).

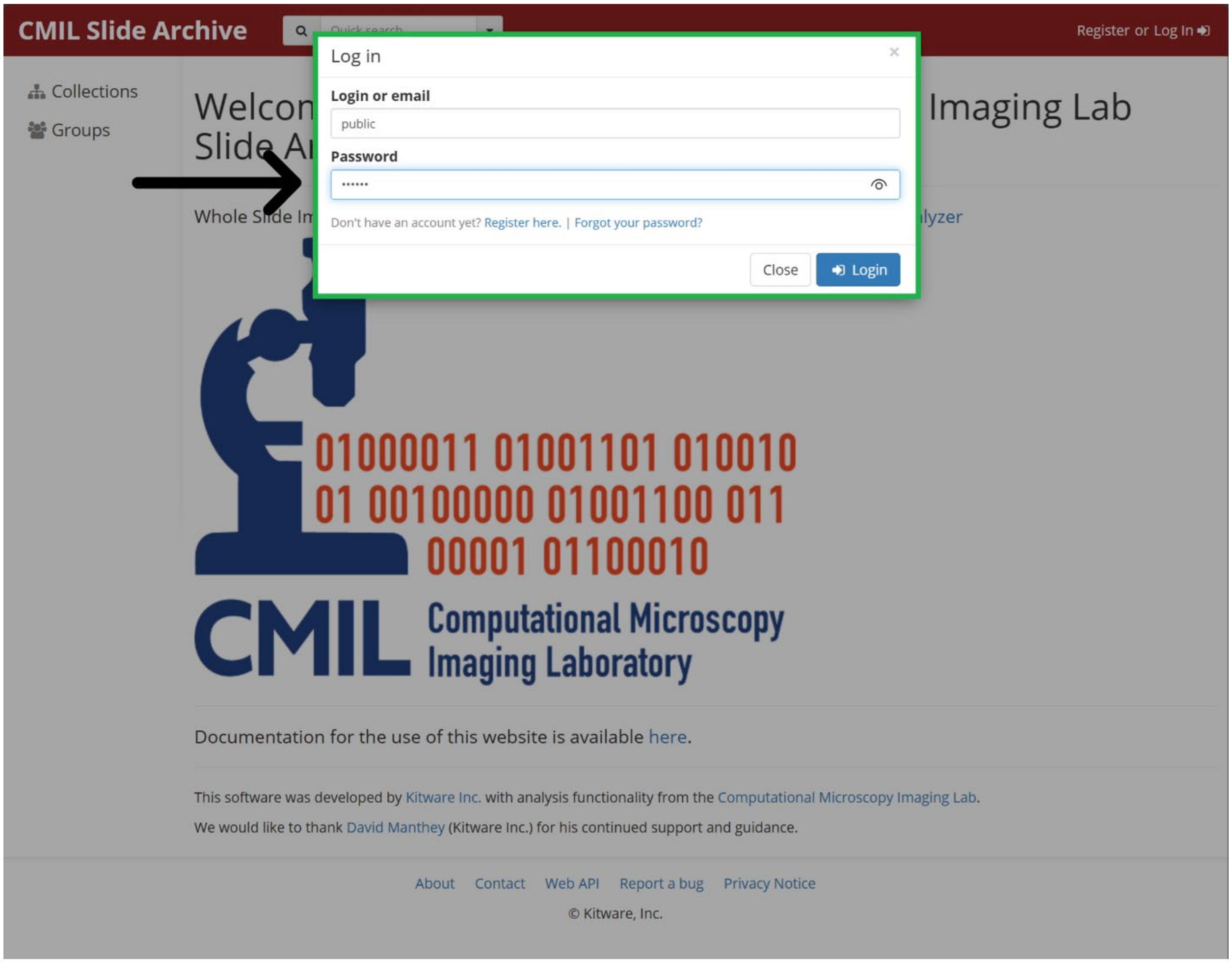


*Supplementary Figure 6:* Login screen to enter Computational Microscopy Imaging Lab Slide Archive using public credentials.

The WSIs used in clinical use-case 1 and 2 are available under Collections/Journals/ WSI_PaulRodrigues_NatureComm_2026 (Supplementary Figure 7).

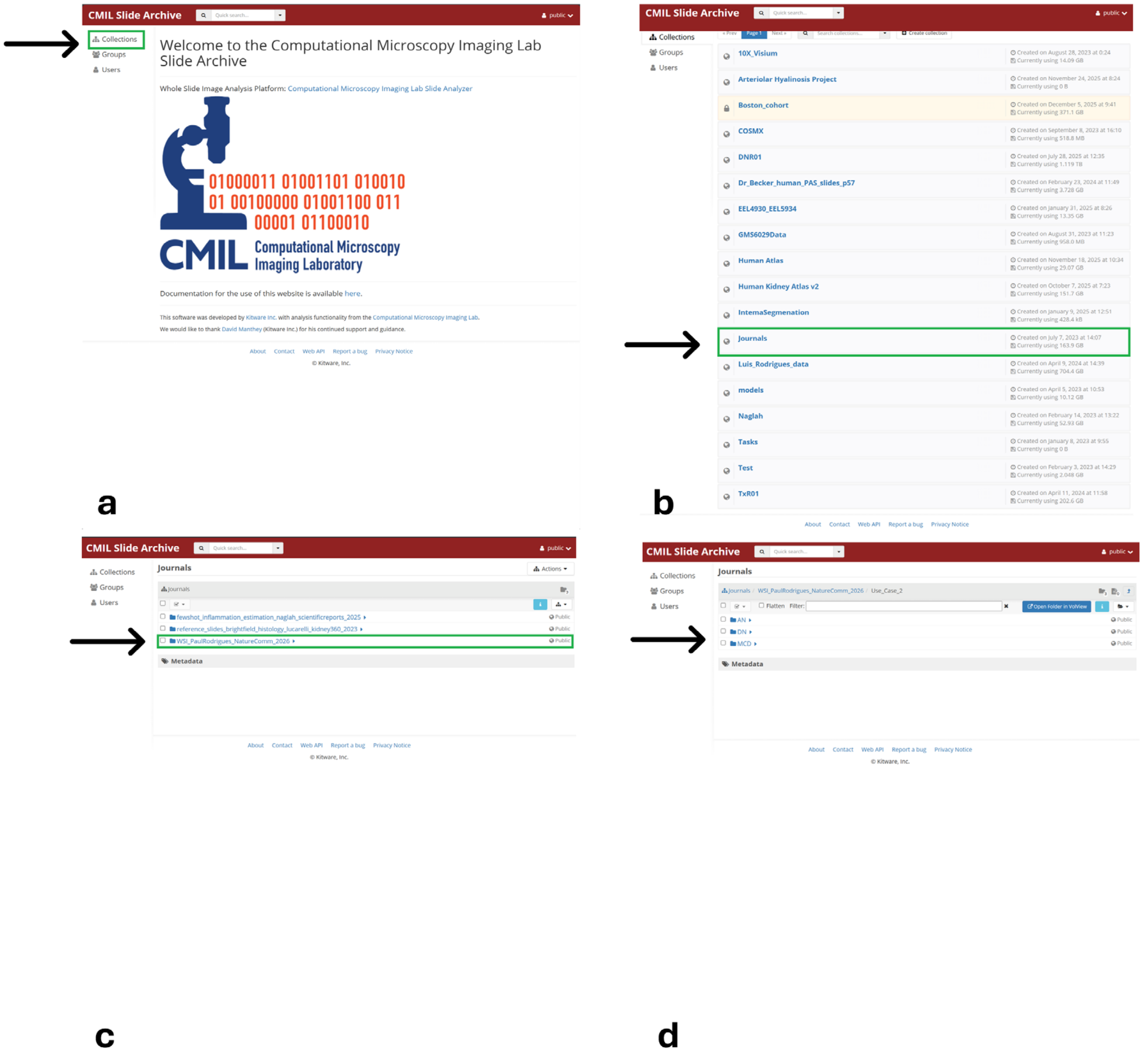


*Supplementary Figure 7*: The WSIs are located at Collections/Journals/WSI_PaulRodrigues_NatureComm_2026. The Use_Case_2 data folder is shown as an example which has 3 distinct disease categories: AN, DN, and MCD. Panels a–d illustrate the sequential steps required to access these WSIs within the Computational Microscopy Imaging Lab Slide Archive.

The user can click on any folder (e.g., AN) to view thumbnail images of all WSIs stored in that folder. Selecting a WSI thumbnail opens a snapshot of the integrated data repository, where AI-generated segmentations, slide metadata, and kidney pathomic features are accessible to the user (Supplementary Figure 8).

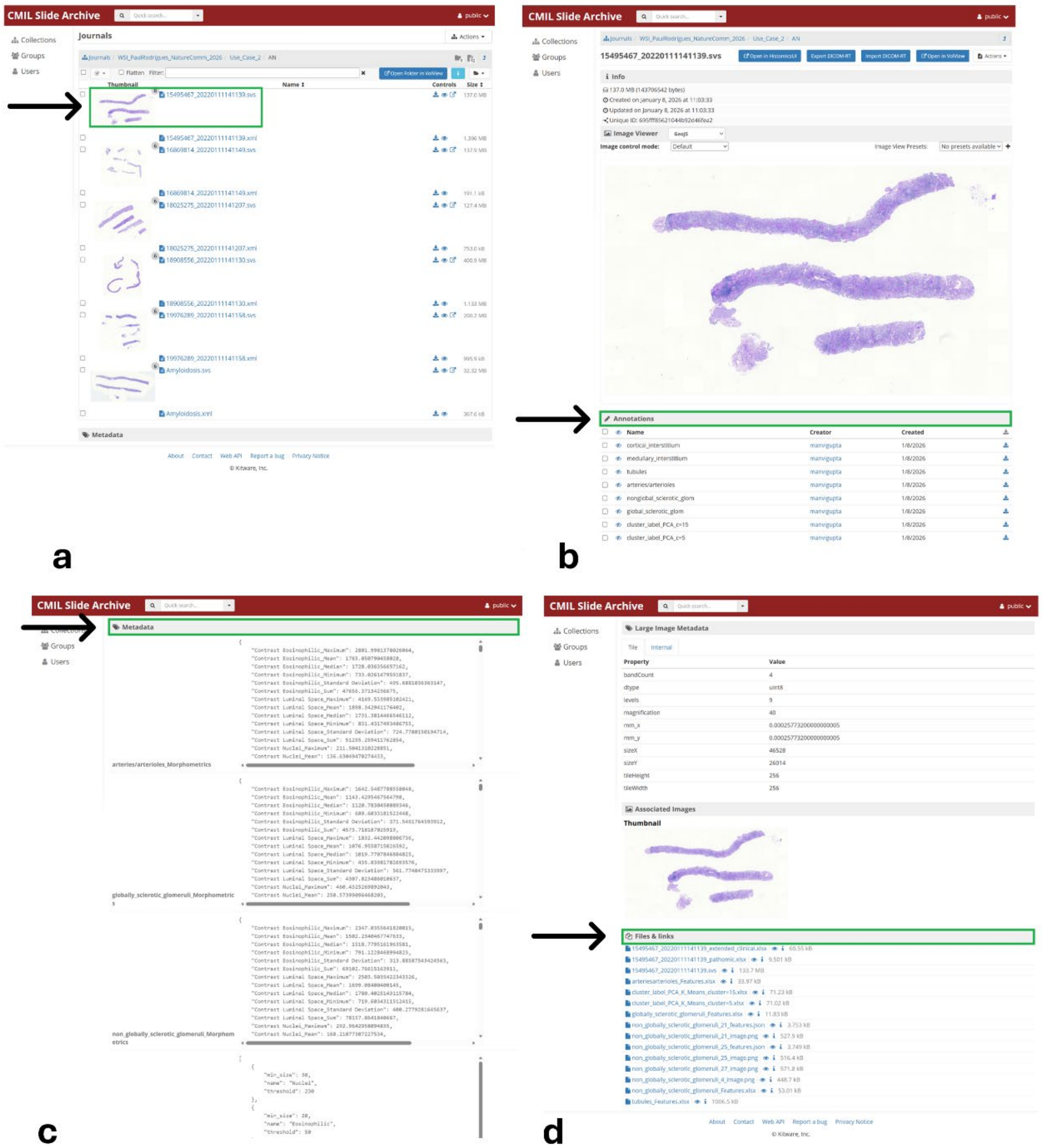

*Supplementary Figure 8*: ComPRePS integrated data repository with WSI, associated AI derived annotations, metadata, morphological and pathomic compartmental features. Panels a–d illustrate the steps required to access this data.

Clicking “Open in HistomicsUI” launches an interactive viewer that supports multi-resolution navigation, zooming, and inspection of annotated tissue structures at cellular detail (Supplementary Figure 9).

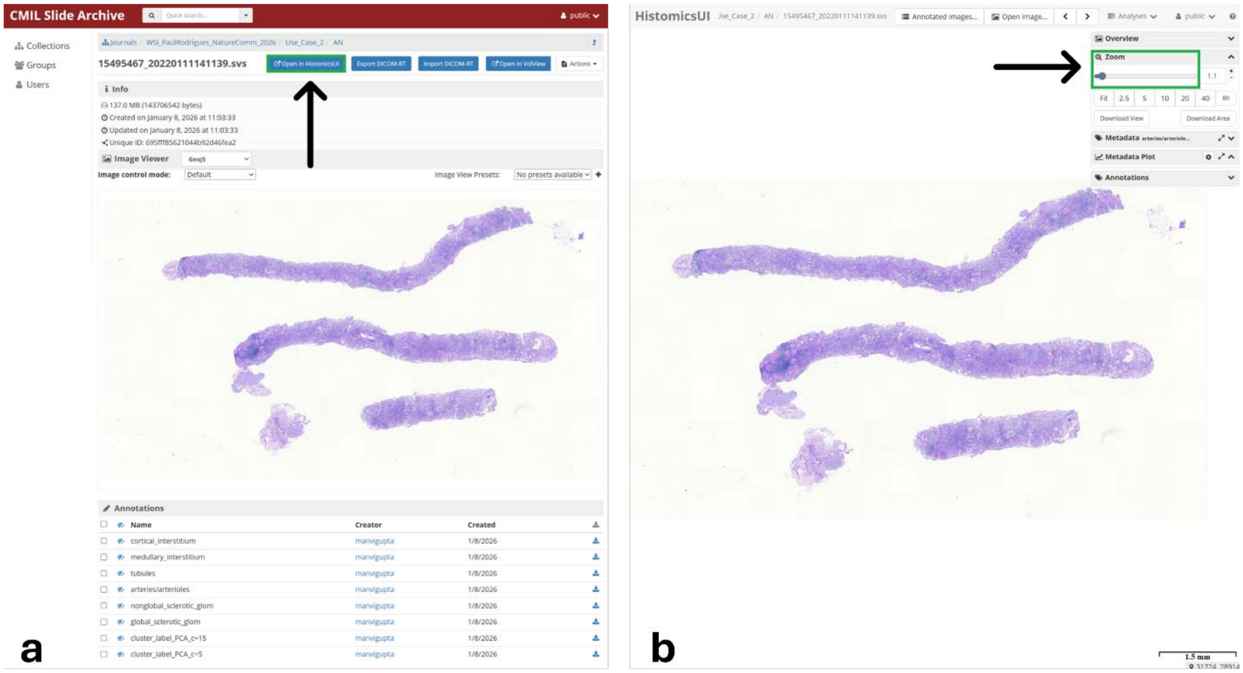


*Supplementary Figure 9*: ComPRePS interactive UI for visualization of tissue compartments and AI segmentations via multi-resolution zooming. Panel a shows the “Open in HistomicsUI” link, which launches the interactive viewer (Panel b).

Upon opening a WSI in HistomicsUI, computational annotations are available under the “Other” section within Annotations and can be toggled on or off using the eye icon (eight JSON-based compartmental annotations relevant to the use-case 2 is shown in Supplementary Figure 10).

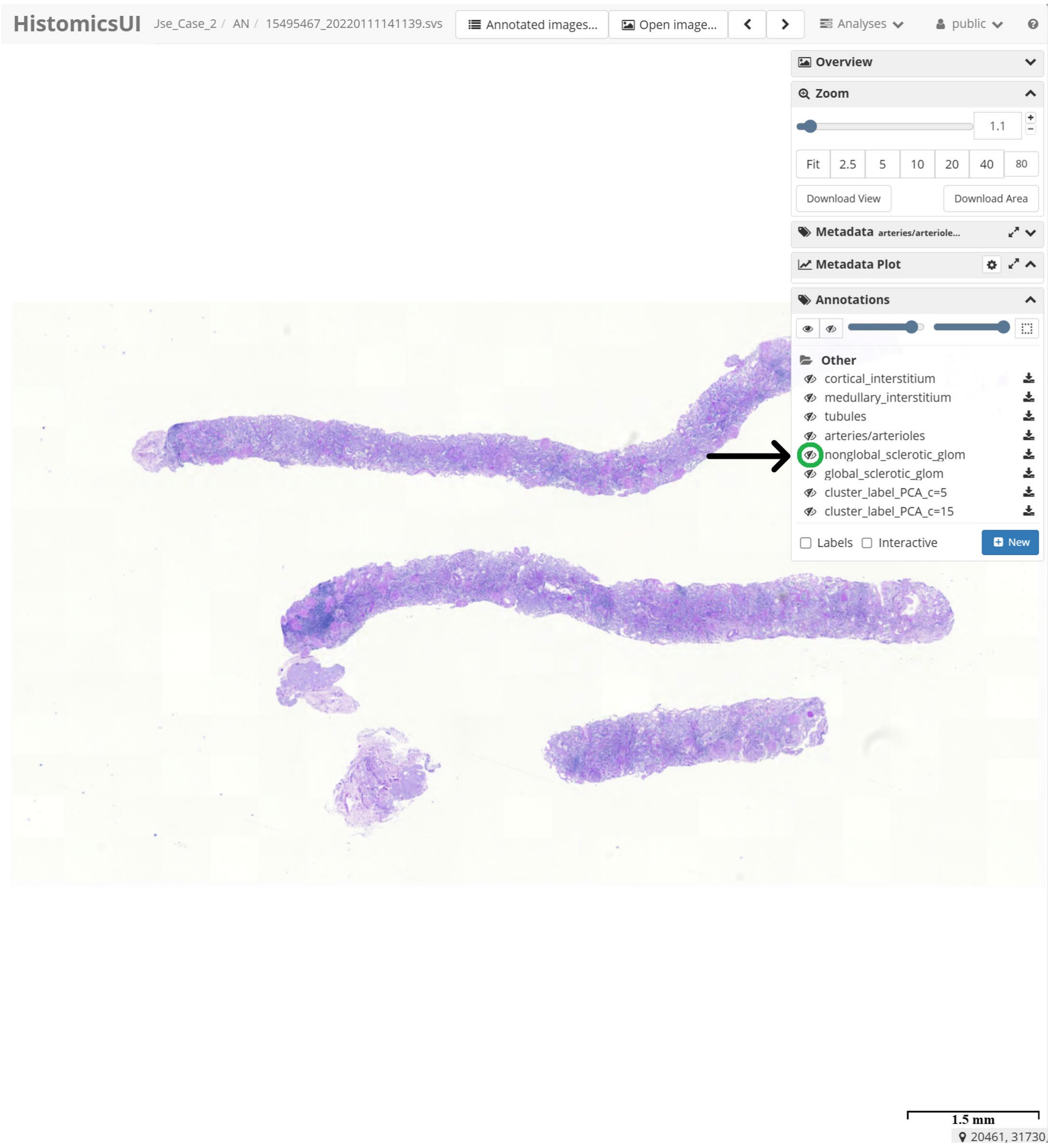


*Supplementary Figure 10*: Visualization of AI segmentations via HistomicsUI. “Annotations” contain multiple AI generated masks and cluster labels performed using K-means clustering.

As shown in Supplementary Figure 10, turning on the eye icon corresponding to each annotation enables overlay of the corresponding boundaries on the WSI. Supplementary Figure 11 illustrates activation of non-globally sclerotic glomeruli (green), tubules (blue), arteries/arterioles (magenta), and globally sclerotic glomeruli (cyan). The segmentation masks can be examined in low-resolution (zoom-out whole slide view) as well as magnified for improved visualization. Due to the large number of cloud-rendered annotations, brief rendering delays may occur depending on the backend server load at that moment. Similarly, other segmented compartments can be visualized in distinct colors.

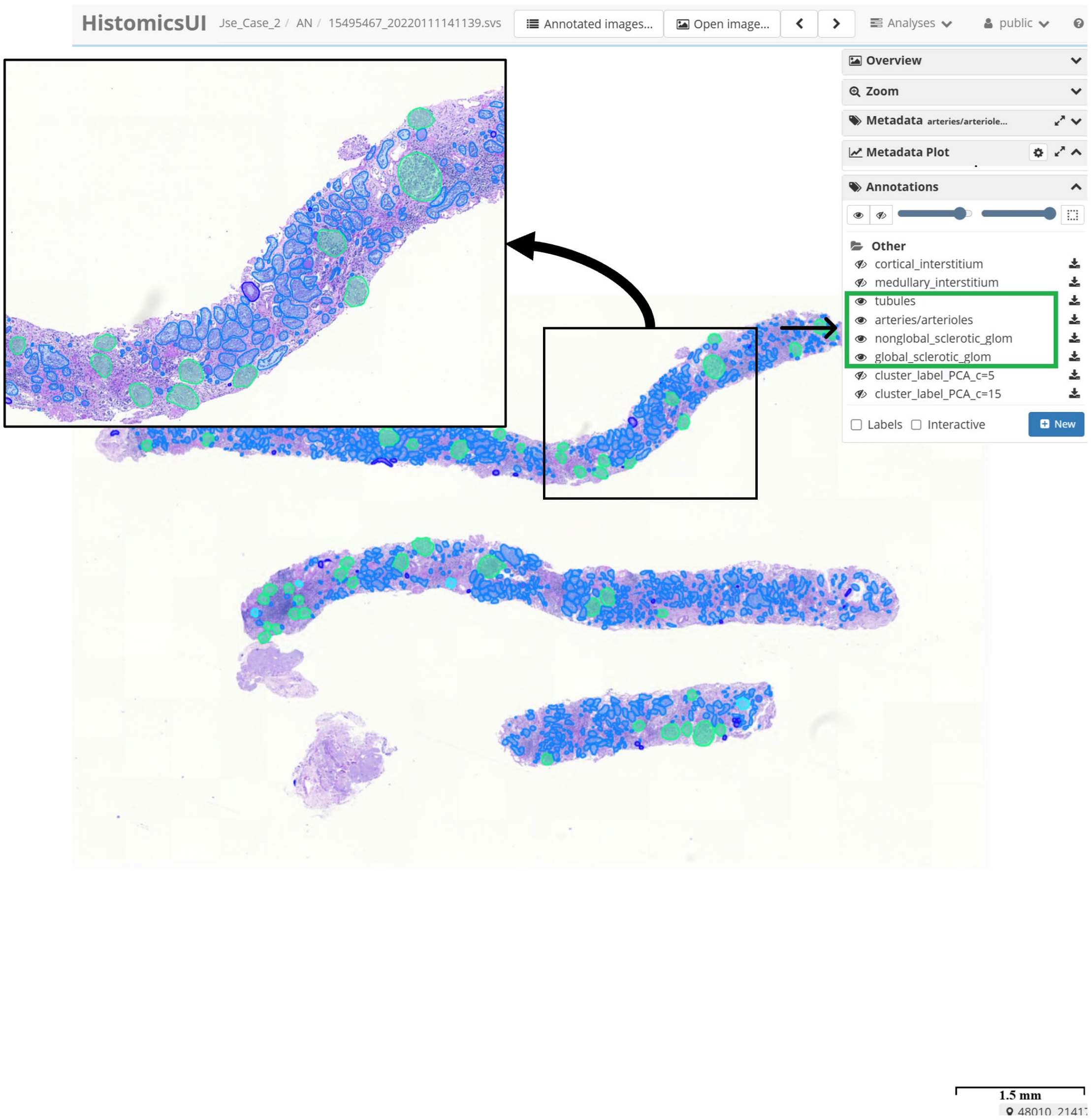


*Supplementary Figure 11*: Compartmental annotations can be toggled in HistomicsUI to overlay segmented renal structures on WSIs for interactive, multi-resolution visualization. The larger black box shows a higher-magnification view of the region outlined by the smaller box.

Segmented masks and corresponding summarized structural measurements (e.g., size and length) for each renal compartment can be downloaded by users in JSON format for local analysis demonstrated in Supplementary Figure 12.

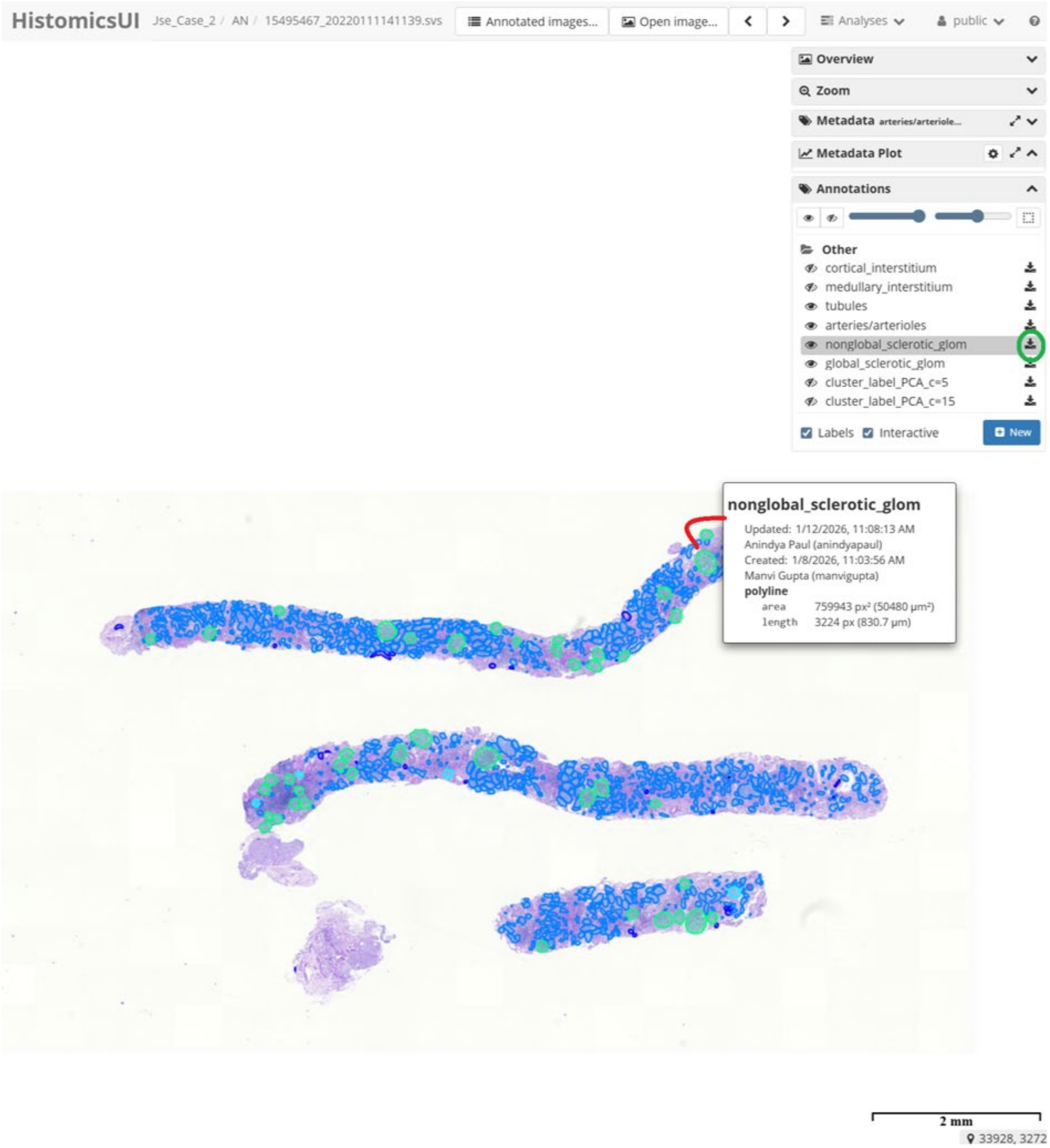


*Supplementary Figure 12*: Download option of segmented structures for porting to other systems or for offline analysis.

#### References


[1] "Multi-compartment segmentation." https://github.com/SarderLab/Multi-Compartment-Segmentation (accessed.

[2] B. G. e. al, "Automated computational detection of interstitial fibrosis, tubular atrophy, and glomerulosclerosis," *Journal of the American Society of Nephrology,* vol. 32, no. 4, pp. 837–850, 2021, doi: https://doi.org/10.1681/asn.2020050652.

[3] Y. Zheng *et al.*, "Deep-Learning–Driven Quantification of Interstitial Fibrosis in Digitized Kidney Biopsies," *The American Journal of Pathology,* vol. 191, no. 8, pp. 1442–1453, 2021/08/01/ 2021, doi: https://doi.org/10.1016/j.ajpath.2021.05.005.

[4] Y. Chen, et al, "Clinical relevance of computationally derived attributes of peritubular capillaries from kidney biopsies," *Kidney360* pp. 648–658, 2023, doi: https://doi.org/10.34067/kid.0000000000000116.

[5] Y. C. e. al, "A whole-slide image (WSI)-based immunohistochemical feature prediction system improves the subtyping of lung cancer," *Lung Cancer,* vol. 165, pp. 18–27, 2022, doi: https://doi.org/10.1016/j.lungcan.2022.01.005.

[6] "PTC Feature Extraction." https://github.com/yijiangchen/PTC_feature_extraction (accessed.

[7] J. B. Kopp *et al.*, "Podocytopathies," *Nature Reviews Disease Primers,* vol. 6, no. 1, p. 68, 2020/08/13 2020, doi: 10.1038/s41572-020-0196-7.

[8] A. Fukuda *et al.*, "Growth-dependent podocyte failure causes glomerulosclerosis," (in eng), *J Am Soc Nephrol,* vol. 23, no. 8, pp. 1351–63, Aug 2012, doi: 10.1681/asn.2012030271.

[9] B. A. Santo, et al, "PodoCount: a robust, fully automated, whole-slide podocyte quantification tool," *Scientific reports* vol. 14, no. 1, pp. 1377–1392, 2024, doi: https://doi.org/10.1016/j.ekir.2022.03.004.

[10] "Intimal Arterial Fibrosis Segmentation." https://github.com/SarderLab/Intimal_Arterial_Fibrosis_Segmentation (accessed.

[11] "sarderlab/histomicstk_extras." https://hub.docker.com/r/sarderlab/histomicstk_extras (accessed.

[12] "Frozen Glom Segmentation." (accessed.

[13] R. K. J. e. al, "Atypical ductal hyperplasia: Interobserver and intraobserver variability," *Modern Pathology,* vol. 24, no. 7, pp. 917–923, 2011, doi: https://doi.org/10.1038/modpathol.2011.66.

[14] N. Lucarelli *et al.*, "Correlating Deep Learning-Based Automated Reference Kidney Histomorphometry with Patient Demographics and Creatinine," (in eng), *Kidney360,* vol. 4, no. 12, pp. 1726–1737, Dec 1 2023, doi: 10.34067/kid.0000000000000299.

[15] S. P. Border, et al, "FUSION: A web-based application for in-depth exploration of multi-omics data with brightfield histology," *Nature Communications* vol. 16, no. 1, p. 8388, 2025, doi: https://doi.org/10.1038/s41467-025-63050-9

[16] A. B, "Cybersecurity in Healthcare: New Threat to Patient Safety," *Cureus* vol. 17, no. 5, 2025, doi: 10.7759/cureus.83614.

[17] P. S. e. al, "Lustre: Building a file system for 1000-node clusters," *Proceedings of the 2003 linux symposium,* pp. 380–386, 2003.
[18] "Lustre (file system)." [Online]. Available: https://en.wikipedia.org/wiki/Lustre_(file_system).
[19] "Globus | go beyond data." https://www.globus.org/.
[20] "Procedures for Regulated Data." https://www.rc.ufl.edu/documentation/procedures/procedures-for-regulated-data/.
[21] X. Yang *et al.*, "A large language model for electronic health records," *npj Digital Medicine,* vol. 5, no. 1, p. 194, 2022/12/26 2022, doi: 10.1038/s41746-022-00742-2.
[22] "A Girder plugin for exposing slicer execution model's CLI modules over the web." https://github.com/girder/slicer_cli_web .
[23] "MONAI." https://github.com/Project-MONAI/model-zoo.
[24] L. McInnes, Healy, J., Saul, N., Großberger, L, "UMAP: Uniform manifold approximation and projection," *Journal of Open-Source Software,* vol. 3, no. 861, 2018.
[25] "Principal component analysis." https://en.wikipedia.org/wiki/Principal_component_analysis.
